# Brain-derived neurotrophic factor contributes to activity-induced muscle pain in male but not female mice

**DOI:** 10.1101/2023.10.31.565022

**Authors:** Kazuhiro Hayashi, Joseph B. Lesnak, Ashley N. Plumb, Adam J. Janowski, Angela F. Smith, Joslyn K. Hill, Kathleen A. Sluka

## Abstract

Activity-induced muscle pain increases release of interleukin-1β (IL-1β) in muscle macrophages and the development of pain is prevented by blockade of IL-1β. Brain derived neurotrophic factor (BDNF) is released from sensory neurons in response to IL-1β and mediates both inflammatory and neuropathic pain. Thus, we hypothesized that metabolites released during fatiguing muscle contractions activate macrophages to release IL-1β, which subsequently activate sensory neurons to secrete BDNF. To test this hypothesis, we used an animal model of activity-induced pain induced by repeated intramuscular acidic saline injections combined with fatiguing muscle contractions. Intrathecal or intramuscular injection of inhibitors of BDNF-Tropomyosin receptor kinase B (TrkB) signaling, ANA-12 or TrkB-Fc, reduced the decrease in muscle withdrawal thresholds in male, but not in female, mice when given before or 24hr after, but not 1 week after induction of the model. BDNF messenger ribonucleic acid (mRNA) was significantly increased in L4–L6 dorsal root ganglion (DRG), but not the spinal dorsal horn or gastrocnemius muscle, 24hr after induction of the model in either male or female mice. No changes in TrkB mRNA or p75 neurotrophin receptor mRNA were observed. BDNF protein expression via immunohistochemistry was significantly increased in L4–L6 spinal dorsal horn and retrogradely labelled muscle afferent DRG neurons, at 24hr after induction of the model in both sexes. In cultured DRG, fatigue metabolites combined with IL-1β significantly increased BDNF expression in both sexes. In summary, fatigue metabolites release, combined with IL-1β, BDNF from primary DRG neurons and contribute to activity-induced muscle pain only in males, while there were no sex differences in the changes in expression observed in BDNF.

## Introduction

Chronic pain costs at least $560–635 billion annually in the U.S., affecting up to 50 million Americans, accounting for roughly 20–40% of the population [1, 2]. Chronic musculoskeletal disorders are one of the most prevalent disorders globally [3]. Chronic musculoskeletal pain can be exacerbated by an acute bout of exercise [4], which creates a barrier to participation in exercise programs aimed at reducing pain severity and improving physical function [5, 6]. To study activity-induced muscle pain, we developed an animal model that combines fatiguing muscle contractions with a non-painful, low-dose muscle insult which results in muscle hyperalgesia [7].

Muscle fatigue releases metabolites, including adenosine triphosphate (ATP), protons, and lactate, which induce the release of the inflammatory cytokine interleukin-1β (IL-1β) from macrophages [8, 9]. IL-1β produces both mechanical and thermal hyperalgesia and is upregulated in inflammatory pain models including our activity-induced pain model [9–12]. Blockade of IL-1 receptors prevents hyperalgesia in inflammatory, neuropathic, and activity-induced pain models [9–12]. Peripheral inflammatory cytokines, including IL-1β, produce their effects by promoting the release of pro-nociceptive mediators including brain derived neurotrophic factor (BDNF) [13, 14]. Sensory neurons express both the IL-1 receptor and BDNF [12, 15], and thus could play a role in activity-induced pain.

BDNF, a member of the neurotrophin family, is expressed in small to medium size sensory neurons, and is upregulated after peripheral inflammation [13–18]. It produces its effects through its receptors, Tropomyosin receptor kinase B (TrkB) and a p75 neurotrophin receptor (p75^NTR^) [13, 14, 16, 19]. BDNF is released from peripheral and central terminals of sensory neurons and likely produces its effects through multiple mechanisms [13, 17, 18, 20–22]. In the spinal cord, BDNF contributes to sensitization of dorsal horn neurons by enhancing NMDA receptor neurotransmission, glial cell activation, and reducing GABA signaling [23–25]. In muscle, BDNF-TrkB plays a role in muscle function, is increased and released from muscle in response to exercise [26] and peripheral BDNF injection decreases pain thresholds [27, 28]. These data suggest that BDNF could produce effects peripherally within the muscle or centrally in the spinal cord to enhance activity-induced pain.

Research on BDNF from sensory neurons is contradicting, showing potential model- and sex-specific roles. Genetic deletion of BDNF from Nav1.8 positive nociceptors prevents inflammatory pain [29], suggesting sensory neuron-derived BDNF is important in the development of hyperalgesia. In contrast, other studies showed genetic deletion of BDNF from sensory neurons reverse pain by spinal nerve ligation in both sexes, but not partial sciatic nerve ligation, paclitaxel, or Complete Freund’s adjuvant in either male or female mice [30, 31]. Genetic depletion showed mixed effects with loss of hyperalgesia to formalin in males but contradicting effects in females [30, 31]. Interestingly, in a hyperalgesic priming model, genetic deletion of BDNF from sensory neurons prevented the acute and later phase development of hyperalgesia, but only in males [31]. These data suggest that there may be model-specific and sex-specific roles for BDNF in pain. The role of BDNF induced by both muscle fatigue and IL-1β from macrophages on the activity-induced pain is currently unknown.

The present study aimed to investigate the role of BDNF from primary sensory neurons in activity-induced muscle pain. We hypothesized that muscle fatigue metabolites activate macrophages to release IL-1β, activating sensory neurons to secrete BDNF.

## Methods

### Animal care

All experiments were approved by the University of Iowa Animal Care and Use Committee and were conducted in accordance with the National Institute of Health’s Guidelines for the Care and Use of Laboratory Animals. A total of 224 male and 224 female C57BL/6J mice (Jackson Laboratories, Bar Harbor, ME, USA), aged 6 to 8 weeks were used in this study. An equal number of male and female mice were used for each experiment. Mice were housed under a 12hr/12hr light/dark cycle with access to food and water *ad libitum*. Mice were randomly assigned to each group using computer generated stratified block randomization. The investigator was blinded to group during behavioral measurements and sample assessment.

### Animal Model of Activity-Induced Muscle Pain

The activity-induced muscle pain model was induced by 2, 20μl acidic pH 5.0 saline injections, separated by 5 days, in the left gastrocnemius muscle combined with 6 min of electrically stimulated fatiguing muscle contractions [7]. Mice were anesthetized with 2–4% isoflurane and given an intramuscular (i.m.) injection into the left gastrocnemius of 20μl normal 0.9% saline (Hospira, Lake Forest, IL, USA) adjusted to pH 5.0 ± 0.1 with HCl. The unbuffered pH 5.0 saline injections reduce muscle pH to approximately 6.9 [32], which is comparable to decreases seen after intense exercise [33–35].

Five days after the first i.m. injection of 20μl pH 5.0 saline, mice were anesthetized with 2–4% isoflurane, needle electrodes were inserted into the belly of the left gastrocnemius muscle followed by 6 min of submaximal fatiguing contractions using a custom-built device. The following parameters were used: a duty cycle of 47% (3.75sec on, 4.25sec off) using trains of stimulations at 40Hz and an amplitude of 7V. This protocol results in approximately 50% reduction in muscle force [7]. Immediately following the fatiguing muscle contractions, the mice were given a second i.m. injection of 20μl pH 5.0 saline into the same gastrocnemius muscle. Pain free control mice received 2, 20μl pH 7.2 ± 0.1 saline injections, separated by 5 days, in the gastrocnemius muscle [7].

### Measurement of Muscle Withdrawal Thresholds

Pain was measured by determining the muscle withdrawal threshold (MWT) of the bilateral gastrocnemius muscles. MWT was measured by applying force-sensitive tweezers to the belly of the gastrocnemius muscle, where lower thresholds indicate greater sensitivity [36]. Prior studies show anesthetizing the skin during this measurement did not change withdrawal thresholds, but anesthetizing the deep tissue increased the threshold, thus validating this as a measurement of muscle hyperalgesia [36].

Briefly, mice were acclimated to the testing paradigm in two 3–5 min sessions over a 2-day period prior to baseline assessment. Mice were placed in a gardener’s glove, the hindlimbs held in extension, and the muscle squeezed with force-sensitive tweezers until the animal withdrew its hindlimb. Five minutes elapsed between each MWT assessment to prevent behavioral sensitization to testing. Each animal was tested bilaterally and the average of three measurements was used to determine withdrawal thresholds for each limb.

### Behavior experiments

To test whether BDNF mediated the development and maintenance of activity-induced muscle pain, selective inhibitors were given intrathecally (5μl volume) or intramuscularly (20μl volume) prior to or after (24hr or 1wk) induction of the activity-induced muscle pain model and were compared to vehicle injections (**Table 1**). For the pre-treatment protocol, intrathecal injections were administered 30 min prior to fatiguing contractions and intramuscular injections were administered 5 min prior to fatiguing contractions [10, 37]. We used two inhibitors of BDNF-TrkB signaling, ANA-12 and TrkB-Fc, and compared them to vehicle [38, 39]. ANA-12 (1.6μg/μl)(Abcam, Cambridge, UK), a BDNF-TrkB signaling antagonist, was dissolved in 10% DMSO and normal saline [38, 39]. Vehicle control for TrkB antagonist was normal saline with 10% DMSO. TrkB-Fc (100ng/μl)(R&D Systems, Minneapolis, MN, USA) was dissolved in normal phosphate buffered saline and used to block the effects of BDNF [38, 39]. Vehicle control for TrkB-Fc was normal phosphate buffered saline. The doses were determined based on data from previous studies [37–40].

**Table 1.** Study procedures (type of inhibitors)

| Inhibitor | Catalogue number, company | Vehicle |
| --- | --- | --- |
| TrkB antagonist (ANA-12) | ab146196, Abcam | 0.9% saline with 10% DMSO |
| TrkB-Fc | 10266-TB-050, R&D Systems | Phosphate buffered saline |
| DMSO, dimethyl sulfoxide |  |  |

For animals treated prior to the induction of the model, MWT was measured before (baseline) and at 24hr, 72hr, and 1wk after the injection of inhibitor or its vehicle. For animals treated after the induction of the model, MWT was measured before and after induction of the model, and 30 min, 2hr, 24hr, 72hr, and 1wk after injection of the inhibitor or its vehicle (n = 4–8 in each sex per group)(total, inhibitor; n = 144; 72 males and 72 females, vehicle control; n = 144; 72 males and 72 females).

### Ribonucleic acid (RNA) extraction from DRG, spinal dorsal horn, or gastrocnemius muscle

For quantitative polymerase chain reaction (qPCR) experiments, bilateral L4–L6 DRG, spinal dorsal horn, and gastrocnemius muscle were collected 2hr, 24hr, and 1wk after induction of the activity-induced muscle pain model (model; n = 8; 4 males and 4 females, pain free control; n = 8; 4 males and 4 females, respectively for each time point). Mice were euthanized with CO_2_ at a rate of 3 L/min for 1 min following cessation of breathing. The tissues were harvested immediately after euthanasia and placed in RNA later (Invitrogen; Thermo Fisher Scientific, Waltham, MA, USA). The L4–L6 DRGs were removed and pooled for the left and right sides of each animal. Total cellular RNA was extracted from the tissues using QIAzol reagent together with RNeasy Mini Kit (Qiagen, Hilden, Germany), and RNA Clean & Concentrator-5 Kit (Zymo Research, Orange, CA, USA) according to the manufacturer’s instructions. Samples of isolated total RNA were stored at -80°C until analysis.

### Immunohistochemistry

A group of animals were used to extract ipsilateral L4–L6 DRG and bilateral spinal dorsal horn of lamina I-II and III-IV to examine protein expression at the 24hr time point after induction of the activity-induced muscle pain model (model; n = 16; 8 males and 8 females, pain free control; n = 16; 8 males and 8 females). Mice were deeply anesthetized (80mg/kg ketamine and 10mg/kg xylazine, intraperitoneal injection) and transcardially perfused with heparinized saline followed by freshly prepared 4% paraformaldehyde in 0.1 M PB. The L4–L6 DRG and spinal dorsal horn were removed, stored in 4% paraformaldehyde in PB overnight, and then stored in 10–30% sucrose (4°C). The tissues were embedded in OCT (Thermo Fisher Scientific, Waltham, MA, USA) in a cryomold (Sakura Finetek USA, Torrance, CA, USA) and frozen at -20°C. Sections from the tissue were cut at 20μm with a cryostat and placed onto slides. Sections were blocked with 5% normal goat serum followed by an Avidin/Biotin blocking kit (Vector Labs, Burlingame, CA, USA) and then incubated overnight at room temperature with the primary antibody, rabbit anti-mouse BDNF (1:250)(Abcam, Cambridge, UK). The sections were rinsed, blocked with 5% normal goat serum, incubated for one hour at room temperature with the secondary antibody, Biotin-Goat Anti rabbit IgG (1:500)(Jackson Immuno Research, West Grove, PA, USA) and incubated for one hour at room temperature with Streptavidin Alexa Fluor™ 568 conjugate (1:500)(Invitrogen; Thermo Fisher Scientific, Waltham, MA, USA). Slides were then cover-slipped with Vectashield (Vector Labs). We used the same procedure carried out in the absence of primary antibody negative control, which did not show any staining in the DRG or spinal dorsal horn. The primary antibody was previously validated using BDNF conditional null mice [29]. Images of sections were taken on a Leica SP8 confocal microscope (Leica Microsystems, Deerfield, IL, USA) in the Central Microscopy Facility at the University of Iowa. Optical density of BDNF staining was analyzed using Image J software (NIH, Bethesda, MD, USA). For each location, 3 sections were quantified and averaged for each animal. All images were taken under the same conditions and stored for analysis. The investigator was blinded to group during staining, image acquisition, and quantification.

A separate group of animals was used to extract ipsilateral L4–L6 DRG to examine BDNF protein expression in retrogradely labelled muscle afferents (model; n = 8; 4 males and 4 females, control; n = 8; 4 males and 4 females). To label muscle afferent neurons, an incision was made in the skin of an anesthetized mouse with 2–4% isoflurane to expose the left gastrocnemius muscle of the leg after shaving. Fast Blue (Polysciences, Warrington, PA, USA), 0.1mg diluted in 20μl of normal saline, was injected into the muscle for retrograde labeling. The incision was closed with silk sutures. Fast Blue containing neurons were identified by blue fluorescence emission on brief exposure of the cells to ultraviolet light. Induction of the pain model was initiated 1 day after Fast blue injection, and muscles were collected 24hr after induction of pain model (7 days after injection of Fast Blue). The ipsilateral L4–L6 DRG sections were stained with the BDNF primary antibody as above. The total number of retrogradely labelled Fast Blue or BDNF positive cells were counted from each animal, 3 sections in each, using Image J software (NIH). The neurons were divided into three size groups (small-sized, <300μm^2^; medium-sized, 300–700μm^2^; large-sized, >700μm^2^) [41]. The small- or medium-sized neurons highly express the neurochemical marker of C-fibers, but not A-fibers, in lumber DRGs of mice [41].

### Animal Model of pH4 pain

To test if the changes of BDNF were unique to the activity-induced pain model, the acidic-saline animal model was induced by 2, 20μl pH 4.0 saline injections, separated by 5 days, in the left gastrocnemius muscle [32, 42]. Mice were anesthetized with 2–4% isoflurane and given i.m. injection into the left gastrocnemius of 20μl normal 0.9% saline (Hospira) adjusted to acidic pH 4.0 ± 0.1 with HCl. Pain free control mice received 2, 20μl pH 7.2 ± 0.1 saline injections, separated by 5 days, in the gastrocnemius muscle (model; n = 16; 8 males and 8 females, pain free control; n = 16; 8 males and 8 females). The bilateral L4–L6 DRG and spinal dorsal horn were collected 24hr after induction of the pH4 pain model, followed by extraction of total cellular RNA as described above.

### Primary DRG Neuronal Culture

To examine the effects of fatigue metabolites and IL-1β on BDNF release from primary DRG neuron, we collected primary DRG neuron as follows [43, 44]. On day 0, naïve mice were euthanized with CO_2_ at a rate of 3L/min for 1 min following cessation of breathing. DRGs from bilateral lumbar regions were removed into DMEM/F12 (Gibco; Thermo Fisher Scientific, Waltham, MA, USA) medium in each animal, and then treated with 1 mg/ml collagenase type 3 (Worthington Biochemical Corporation, Lakewood, NJ, USA) at 37°C for 30 min. Neurons were incubated with Trypsin-EDTA (0.25%)(Gibco; Thermo Fisher Scientific) at 37°C for 30 min. Next, neurons were centrifuged at 300 × g for 5 min, re-suspended in DMEM/F12 medium, and repeated three times. Neurons were manually triturated with a transfer pipette. The dissociated cells were suspended in DMEM/F12 medium containing 10% fetal bovine serum (Gibco; Thermo Fisher Scientific), 100μg/ml penicillin-streptomycin (Gibco; Thermo Fisher Scientific), and 1mM sodium pyruvate (Gibco; Thermo Fisher Scientific) and plated in 24-well plates (Nest Biotechnology, Wuxi, China) containing 50μg/ml poly-D-Lysine (Gibco; Thermo Fisher Scientific) coated for 24hr (37°C, 5% CO_2_).

On day 1, cells were treated with pH 7.4 media (26mM NaHCO_3_ (Research Products International, Mount Prospect, IL, USA), 25mM 4-(2-hydroxyethyl)-1-piperazineethanesulfonic acid (HEPES)(Research Products International) or acidic pH 6.5 media (26mM NaHCO_3_, 20mM 2-(N-morpholino) ethanesulfonic acid (MES)(Millipore Sigma, St. Louis, MO, USA), 5mM HEPES) with or without 1mM ATP (Millipore Sigma)) with or without recombinant IL-1β (50ng/ml)(Invitrogen; Thermo Fisher Scientific, Waltham, MA, USA) for 24hr [45, 46] (n = 8; 4 males and 4 females, respectively for each group).

At the end of the experiments on day 2, the cells were collected for protein analysis and qPCR. Total cellular RNA and protein were extracted using Trizol reagent (Invitrogen; Thermo Fisher Scientific) and RNA Clean & Concentrator-5 Kit (Zymo Research, Orange, CA, USA) according to the manufacturer’s instructions. Samples of isolated total cellular RNA and protein were stored at -80°C until analysis.

### Quantitative polymerase chain reaction (qPCR)

Total RNA concentrations were measured on a NanoDrop One (Thermo Fisher Scientific). The ratio of A260/A280 was greater than 1.8 for all samples. Total RNA of 100ng was subjected to reverse transcription using the AffinityScript qPCR cDNA Synthesis Kit (Agilent Technologies, Santa Clara, CA, USA), according to the manufacturer’s instructions. The synthesized cDNA was stored at -20°C, and then used as a template for qPCR with the Power SYBR Green PCR Master Mix (Applied Biosystems; Thermo Fisher Scientific, Waltham, MA, USA).

All gene-specific PCR primers were obtained from Primer Bank and manufactured by Integrated DNA Technologies (Coralville, IA, USA) [47–49]. All qPCR primers were designed and assessed for specificity of target gene using the Primer-BLAST software (National Center for Biotechnology Information, National Library of Medicine, Bethesda, MD, USA; http://www.ncbi.nlm.nih.gov/tools/primer-blast/). Forward and reverse primer sequences for 36B4, BDNF, TrkB, and p75^NTR^ are shown in **Table 2**. Due to high expression levels, 36B4 was used as an endogenous control to normalize results for each sample according to our preliminary results. The qPCR analysis was performed with a QuantStudio 7 Flex Real-Time PCR System (Applied Biosystems; Thermo Fisher Scientific) using the ΔΔCt method [50]. The qPCR was performed as follows: 2 min at 50°C, 10 min at 95°C, 40 cycles of 15 sec at 95°C and 1 min at 60°C. The qPCR analysis was run in triplicate with cycle threshold values averaged. All analyses were done with the experimenter blinded to group.

**Table 2.**
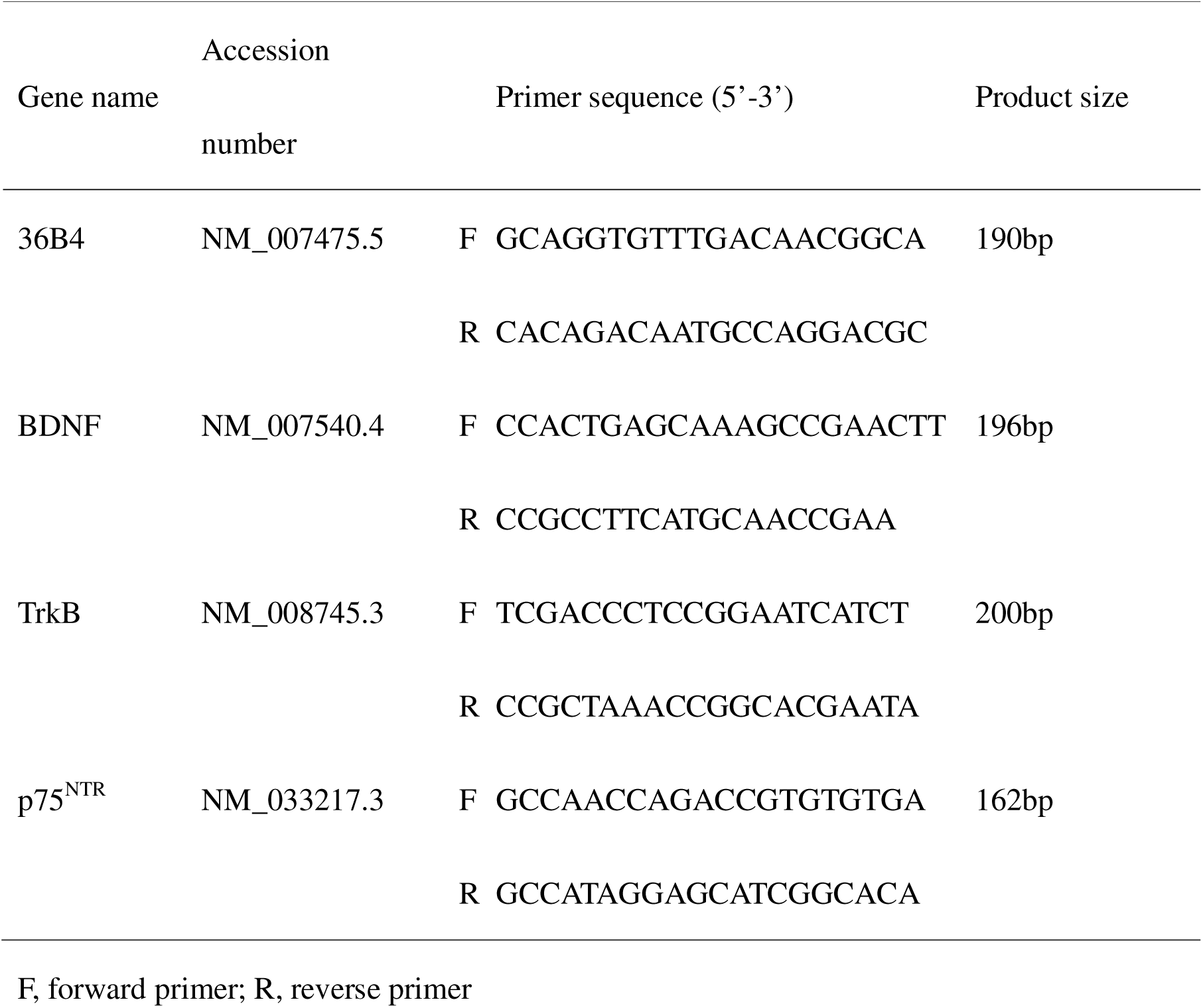
Study procedures (primer sets used in quantitative polymerase chain reaction)

### Enzyme-linked immunosorbent assay (ELISA)

The concentrations of protein expression of BDNF were measured from cultured DRG. The protein concentrations were analyzed using a Quick Start Bradford Protein Assay Kit with bovine serum albumin as a standard (Bio-Rad Laboratories, Hercules, CA, USA). The samples were standardized by diluting in the diluent. ELISA kits (Mouse BDNF ELISA Kit PicoKine™ EK0309)(Boster Biological Technology, Pleasanton, CA, USA) were processed using standard procedures provided by the manufacturer. The absorbance was measured using a Cytation 5 (Biotek, Shoreline, WA, USA) at 450nm. Two standards were used to generate a standard curve across the detectable range (R^2^>0.99). Samples were run in duplicate and coefficients of variation were assessed. All samples showed a coefficient of variation less than 20%. All analyses were done with the experimenter blinded to group.

### Experimental design

The main goal of this study was to examine if up-regulation of BDNF contributed to activity-induced muscle pain. (1) To test whether BDNF mediated activity-induced muscle pain, selective inhibitors of BDNF and its receptors were given prior to or after induction of the activity-induced muscle pain model in muscle or spinal cord, and compared to vehicle injections. MWT was measured before and after injection of BDNF receptor inhibitors for 1 week. (2) To test whether the activity-induced pain model produced changes in expression of BDNF mRNA, BDNF mRNA was measured in DRG, spinal dorsal horn, and gastrocnemius muscle 2hr, 24hr, and 1wk after induction of the model and compared to controls. (3) To test if there were changes in expression of BDNF protein in DRG, we examined immunohistochemical stains for BDNF 24hr after induction of the model in the DRG and spinal cord and compared to controls. (4) To test whether the changes in BDNF protein were localized to muscle afferents, we injected Fast blue into the muscle to retrogradely label muscle DRG neurons. We examined their colocalization with BDNF 24hr after induction of the model. (5) To test if fatigue metabolites combined with IL-1β increase BDNF production, we treated cultured DRG with a combination of acidic pH, ATP, and IL-1β. We then measured mRNA for BDNF and its receptors TrkB and p75^NTR^ in DRG, as well as BDNF protein using an ELISA.

### Statistical analysis

For measurement of MWT, power analysis determined the minimal sample size for withdrawal thresholds was 8 mice per group at 80% power and p<0.05, comparing means for two independent groups (mean 1, 98%; mean 2, 60%; SD1, 25; SD2, 25) [9, 10]. For qPCR and protein analysis, we used the effect size from preliminary results (mean 1, 3,000; mean 2, 300; SD1, 2,450; SD2, 10) to estimate a sample size of 8 per group.

All data are reported as mean ± S.E.M. Group (inhibitor and vehicle control) and differences in MWT for both the ipsilateral and contralateral MWT were analyzed with a repeated measures ANOVA with fixed effects for group and sex, followed by post hoc Tukey’s test. The quantification of qPCR and Immunohistochemistry compared two groups (model and pain free control) with a two-way ANOVA, followed by post hoc testing using an unpaired *t* test with a Bonferroni correction. The quantification of qPCR and ELISA of cultured DRG were analyzed with a one-way ANOVA, followed by post hoc testing using unpaired *t* tests with a Bonferroni correction. A Cohen’s d effect size was calculated. All data were analyzed using IBM SPSS Statistics 28.0 software (IBM, Armonk, NY, USA). A p-value of less than 0.05 was considered statistically significant.

## Results

### Inhibition of BDNF in the activity-induced pain model reduces muscle hyperalgesia in male but not female mice

As previously reported [7, 51], induction of the activity-induced pain model produced a sex-dependent muscle pain phenotype. Females demonstrated a bilateral decrease in MWT while males showed a unilateral decrease in MWT.

To test whether BDNF mediated activity-induced muscle pain, selective inhibitors of BDNF receptors were given prior to or after induction of the activity-induced muscle pain model and compared to vehicle injections. Intrathecal or intramuscular injection of inhibitors of BDNF-TrkB signaling given 24hr after induction of the activity-induced pain model temporarily alleviated the decrease in MWT in male, but not in female mice, when compared to vehicle controls (**Fig. 1**). Significant effects for group and sex occurred for both inhibitors (ANA-12 and TrkB-Fc) and both routes of injection (intrathecally and intramuscularly) (**Table 3**). There were significant increases in MWT in males, but not females, 30 min after intrathecal ANA-12 (p=0.03*, d=1.80), intrathecal TrkB-Fc (p=0.04*, d=1.12), intramuscular ANA-12 (p=0.01*, d=1.72), or intramuscular TrkB-Fc (p=0.002*, d=2.31). There were also significant increases in MWT in males, but not females, 2hr after intrathecal ANA-12 (p=0.01*, d=1.45), or intrathecal TrkB-Fc (p=0.01*, d=1.46). The effects of the BDNF inhibitors on MWT returned to baseline levels 24hr after injection of the BDNF inhibitors.

**Fig. 1.**
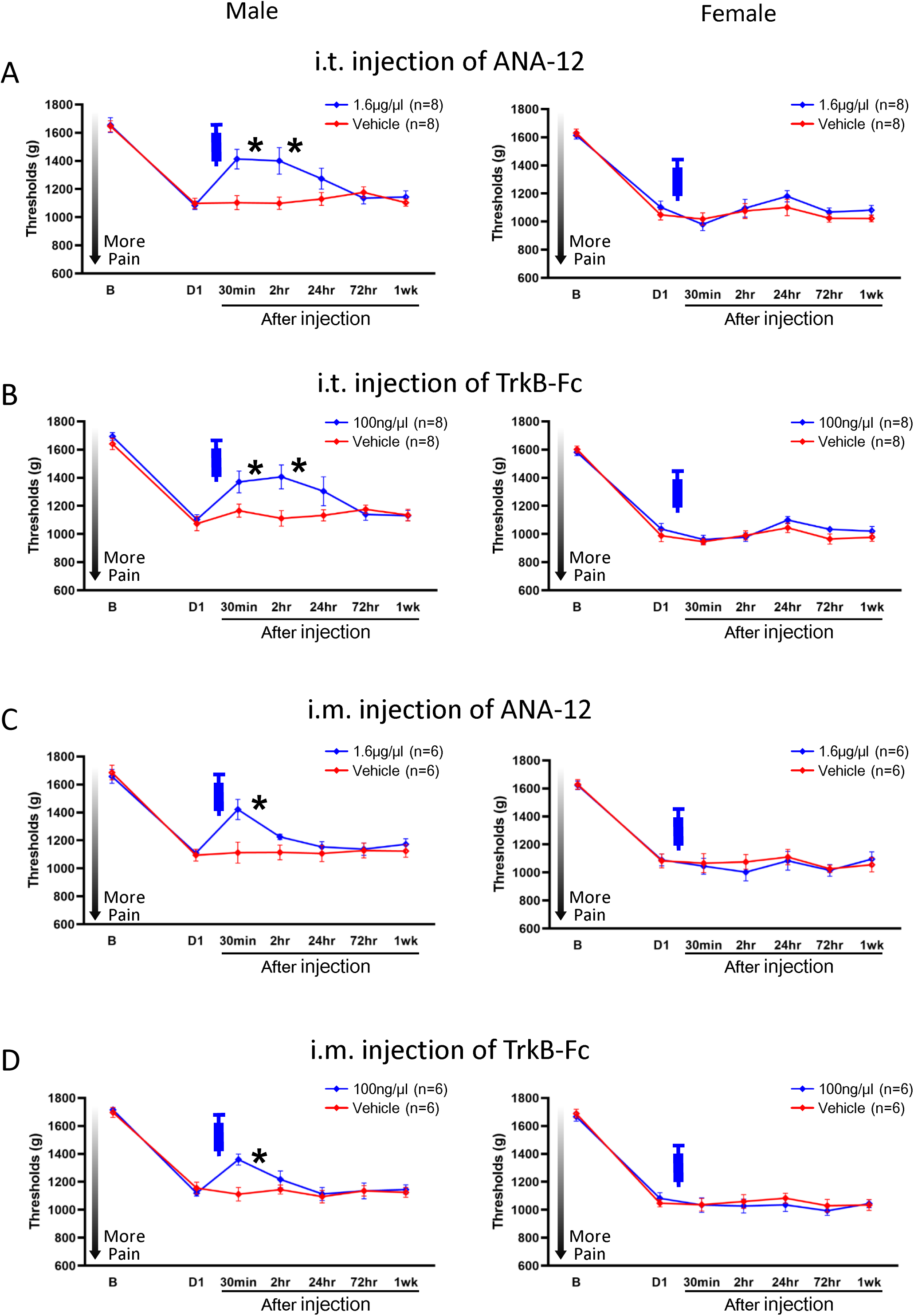
Ipsilateral muscle withdrawal thresholds with pharmacological inhibition intrathecally or intramuscularly 24hr after induction of the model. The decrease in muscle withdrawal thresholds in male, but not female, mice was alleviated by administration of BDNF inhibitors intrathecally or intramuscularly 24hr after induction of the model (A–D). Data is reported as mean ± S.E.M. *, versus vehicle control; p<0.05. i.m., intramuscularly; i.t., intrathecally.

**Table 3.** Statistical results (results for repeated-measures analysis of variance for muscle withdrawal threshold ipsilaterally).

|  | Effect for Group | Effect for Sex | Effect for Group*Sex |
| --- | --- | --- | --- |
| *24hr after induction of the activity-induced muscle pain model* |  |  |  |
| i.t. injection of ANA-12 | $F_{1,28}=5.47$; $p=0.02^*$ | $F_{1,28}=11.49$; $p=0.002^*$ | $F_{1,28}=2.28$; $p=0.14$ |
| i.t. injection of TrkB-Fc | $F_{1,28}=4.93$; $p=0.03^*$ | $F_{1,28}=32.52$; $p<0.001^*$ | $F_{1,28}=1.61$; $p=0.21$ |
| i.m. injection of ANA-12 | $F_{1,20}=1.06$; $p=0.31$ | $F_{1,20}=7.11$; $p=0.01^*$ | $F_{1,20}=2.18$; $p=0.15$ |
| i.m. injection of TrkB-Fc | $F_{1,20}=0.54$; $p=0.46$ | $F_{1,20}=12.49$; $p=0.002^*$ | $F_{1,20}=1.92$; $p=0.18$ |
| *Prior to induction of the activity-induced muscle pain model* |  |  |  |
| i.t. injection of ANA-12 | $F_{1,28}=2.85$; $p=0.10$ | $F_{1,28}=15.58$; $p<0.001^*$ | $F_{1,28}=1.21$; $p=0.28$ |
| i.t. injection of TrkB-Fc | $F_{1,28}=1.40$; $p=0.24$ | $F_{1,28}=28.24$; $p<0.001^*$ | $F_{1,28}=2.70$; $p=0.11$ |
| i.m. injection of ANA-12 | $F_{1,20}=0.006$; $p=0.93$ | $F_{1,20}=4.24$; $p=0.05$ | $F_{1,20}=0.11$; $p=0.73$ |
| i.m. injection of TrkB-Fc | $F_{1,20}=0.64$; $p=0.43$ | $F_{1,20}=24.68$; $p<0.001^*$ | $F_{1,20}=0.91$; $p=0.35$ |

*1wk after induction of the activity-induced muscle pain model*

|  |  |  |  |
| --- | --- | --- | --- |
| i.t. injection of ANA-12 | $F_{1,12}=0.002$; $p=0.96$ | $F_{1,12}=9.08$; $p=0.01^*$ | $F_{1,12}=0.54$; $p=0.47$ |
| i.t. injection of TrkB-Fc | $F_{1,12}=0.32$; $p=0.57$ | $F_{1,12}=10.13$; $p=0.008^*$ | $F_{1,12}=0.004$; $p=0.95$ |
| i.m. injection of ANA-12 | $F_{1,12}=0.01$; $p=0.90$ | $F_{1,12}=1.18$; $p=0.29$ | $F_{1,12}=0.03$; $p=0.85$ |
| i.m. injection of TrkB-Fc | $F_{1,12}=0.002$; $p=0.96$ | $F_{1,12}=2.40$; $p=0.14$ | $F_{1,12}=0.07$; $p=0.79$ |

\*, $p<0.05$ with a repeated measures ANOVA.

Intrathecal injection of the BDNF inhibitors prior to induction of the activity-induced muscle pain model reduced the decrease in MWT ipsilaterally in male mice, but not intramuscular injection, when compared to vehicle controls (**Fig. 2**). Injection of the inhibitors either intrathecally or intramuscularly prior to induction of the activity-induced muscle pain model had no effect in female mice. Significant effects for group and sex occurred for both inhibitors (ANA-12 and TrkB-Fc) of intrathecal injection (**Table 3**). In males, but not females, the decrease of MWT caused by the pain model was reduced by either ANA-12 intrathecally (p=0.03*, d=1.16), or TrkB-Fc intrathecally (p=0.009*, d=1.51) 24hr after injection of the BDNF inhibitors. However, the effects returned by Day 3 (72hr) after injection.

**Fig. 2.**
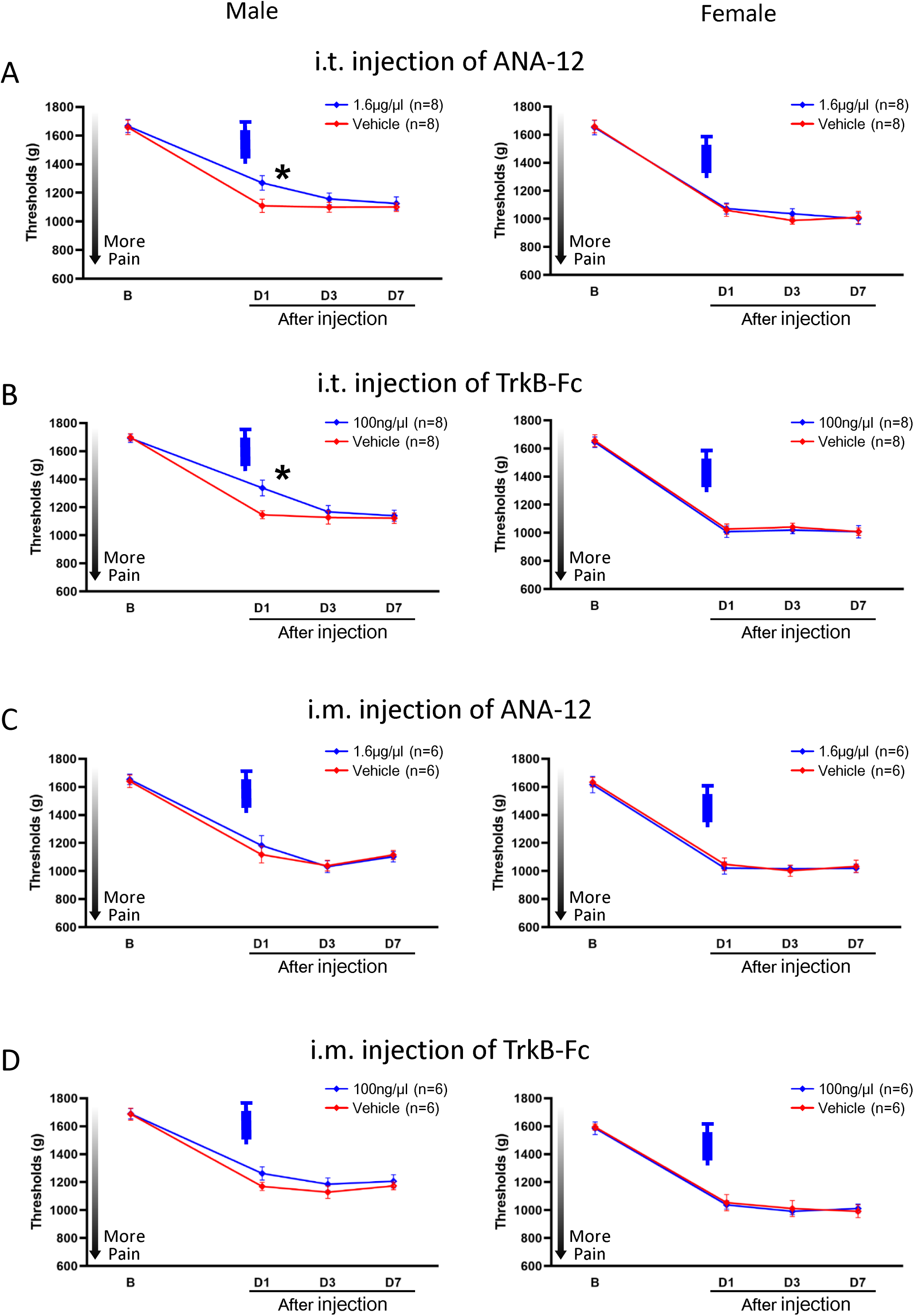
Ipsilateral muscle withdrawal thresholds with pharmacological inhibition intrathecally or intramuscularly prior to induction of the model. The decrease in muscle withdrawal thresholds in male, but not female, mice was reduced by administration of BDNF inhibitors intrathecally, but not intramuscularly, prior to induction of the model (A–D). Data is reported as mean ± S.E.M. *, versus vehicle control; p<0.05. i.m., intramuscularly; i.t., intrathecally.

Neither intrathecal nor intramuscular injection of the BDNF inhibitors, 1wk after induction of the activity-induced pain model, had any effect on the ipsilateral side in male or female mice **(Supplemental Fig. 1)(Table 3**). Injection of the inhibitors either intrathecally or intramuscularly had no effect on the contralateral side in male or female mice at any time periods **(Supplemental Fig. 2–4)(Table 4**). This demonstrates that BDNF is important in the early phase for maintaining hyperalgesia in males in the activity-induced muscle pain model.

**Table 4.** Statistical results (results for repeated-measures analysis of variance for muscle withdrawal threshold contralaterally).

|  | Effect for Group | Effect for Sex | Effect for Group*Sex |
| --- | --- | --- | --- |
| *24hr after induction of the activity-induced muscle pain model* |  |  |  |
| i.t. injection of ANA-12 | $F_{1,28}=2.25$; $p=0.14$ | $F_{1,28}=347.31$; $p<0.001^*$ | $F_{1,28}=0.02$; $p=0.87$ |
| i.t. injection of TrkB-Fc | $F_{1,28}=0.96$; $p=0.33$ | $F_{1,28}=395.12$; $p<0.001^*$ | $F_{1,28}=0.08$; $p=0.77$ |
| i.m. injection of ANA-12 | $F_{1,20}=0.001$; $p=0.98$ | $F_{1,20}=162.88$; $p<0.001^*$ | $F_{1,20}=0.21$; $p=0.64$ |
| i.m. injection of TrkB-Fc | $F_{1,20}=0.02$; $p=0.88$ | $F_{1,20}=171.35$; $p<0.001^*$ | $F_{1,20}=0.06$; $p=0.80$ |
| *Prior to induction of the activity-induced muscle pain model* |  |  |  |
| i.t. injection of ANA-12 | $F_{1,28}=0.04$; $p=0.83$ | $F_{1,28}=331.15$; $p<0.001^*$ | $F_{1,28}=0.003$; $p=0.95$ |
| i.t. injection of TrkB-Fc | $F_{1,28}=0.15$; $p=0.69$ | $F_{1,28}=285.30$; $p<0.001^*$ | $F_{1,28}=0.15$; $p=0.69$ |
| i.m. injection of ANA-12 | $F_{1,20}=0.28$; $p=0.60$ | $F_{1,20}=323.76$; $p<0.001^*$ | $F_{1,20}=0.13$; $p=0.71$ |
| i.m. injection of TrkB-Fc | $F_{1,20}=1.07$; $p=0.31$ | $F_{1,20}=220.79$; $p<0.001^*$ | $F_{1,20}=0.15$; $p=0.70$ |

*1wk after induction of the activity-induced muscle pain model*

|  |  |  |  |
| --- | --- | --- | --- |
| i.t. injection of ANA-12 | $F_{1,12}=0.004$; $p=0.95$ | $F_{1,12}=271.85$; $p<0.001^*$ | $F_{1,12}=0.58$; $p=0.45$ |
| i.t. injection of TrkB-Fc | $F_{1,12}=0.85$; $p=0.37$ | $F_{1,12}=817.67$; $p<0.001^*$ | $F_{1,12}=0.49$; $p=0.49$ |
| i.m. injection of ANA-12 | $F_{1,12}=0.30$; $p=0.59$ | $F_{1,12}=274.79$; $p<0.001^*$ | $F_{1,12}<0.001$; $p=0.99$ |
| i.m. injection of TrkB-Fc | $F_{1,12}=0.14$; $p=0.71$ | $F_{1,12}=166.45$; $p<0.001^*$ | $F_{1,12}=0.04$; $p=0.82$ |

\*, $p<0.05$ with a repeated measures ANOVA.
i.m., intramuscularly; i.t., intrathecally.

### Increased expression of BDNF after induction of the activity-induced muscle pain model

To test whether the activity-induced pain model changed expression of BDNF or its receptors, mRNA was measured in DRG, spinal dorsal horn, and gastrocnemius muscle. There were no sex differences in the expression of mRNA or protein (**Table 5–7**). BDNF mRNA was significantly increased when compared to pain free controls in the ipsilateral L4–L6 DRG (p<0.001*, d=3.39) 24hr after induction of the model, but not at 2hr or 1wk (**Fig. 3**)(**Table 5**). No changes in BDNF mRNA were observed in the contralateral L4–L6 DRG, both ipsilateral and contralateral spinal dorsal horn, or the gastrocnemius muscle 24hr after induction of the model. No changes in TrkB mRNA or p75^NTR^ mRNA were observed in any tissues **(Supplemental Fig. 5, 6)(Table 5**). To test if the changes were unique to the activity-induced pain model, we examined mRNA expression of BDNF in the acidic-saline model induced by 2 i.m. injections of pH 4.0 saline. No changes in BDNF mRNA were observed 24hr after induction of the acidic-saline model when compared to pain free controls in either the ipsilateral or contralateral L4–L6 DRG or spinal dorsal horn **(Supplemental Fig. 7)(Table 8**).

**Fig. 3.**
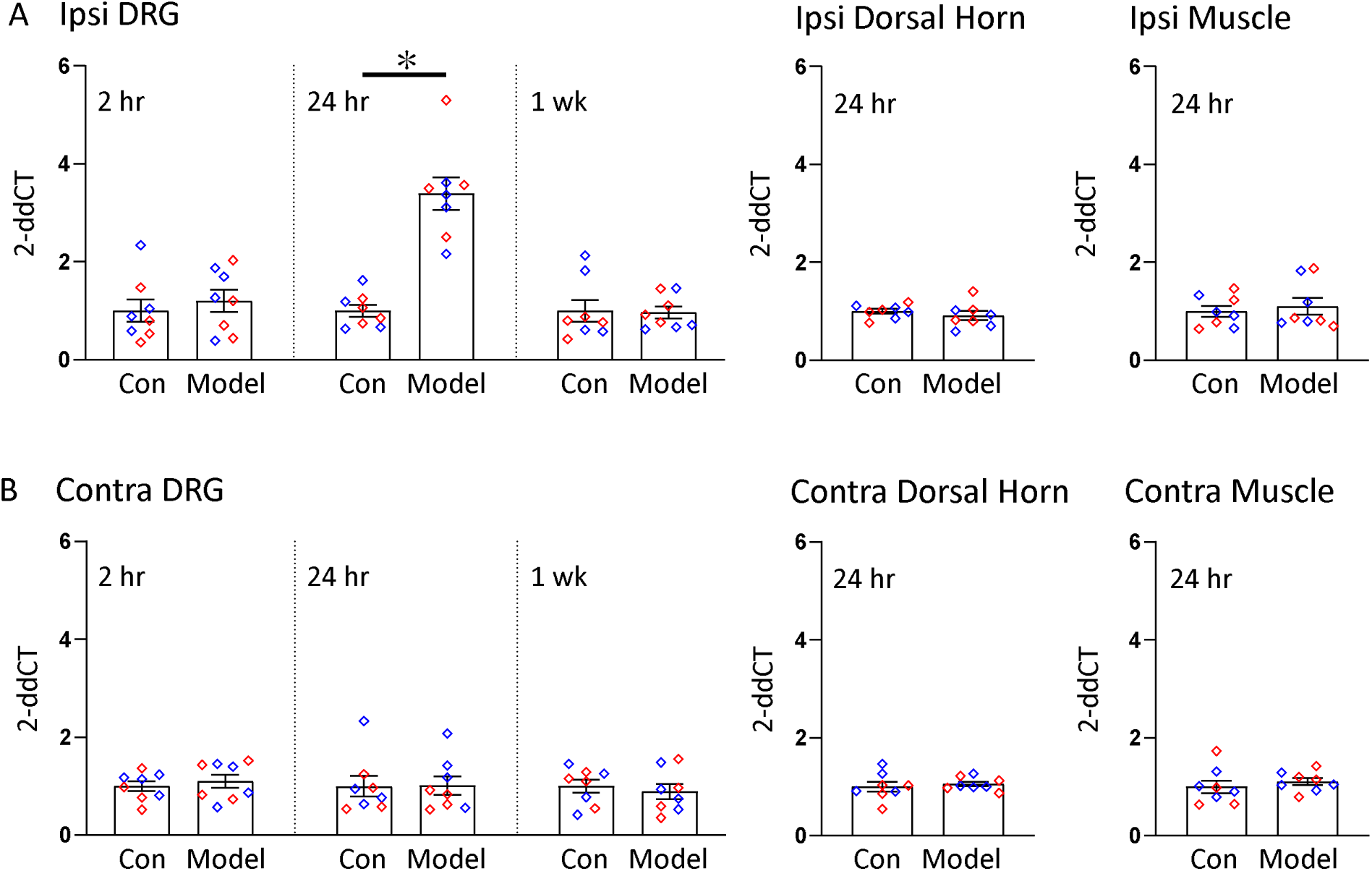
BDNF mRNA expressions after induction of the activity-induced muscle pain. BDNF mRNA in ipsilateral L4–L6 DRG 24hr after induction of the model was significantly increased (A), but not contralateral L4–L6 DRG (B), or either ipsilateral or contralateral spinal dorsal horn or gastrocnemius muscle, when compared to pain free controls. Blue plots represent male data, Red plots represent female data. Data is reported as mean ± S.E.M. N=8 in each group. *, p<0.05. BDNF, brain-derived neurotrophic factor; DRG, dorsal root ganglion; mRNA, messenger ribonucleic acid.

**Table 5.**
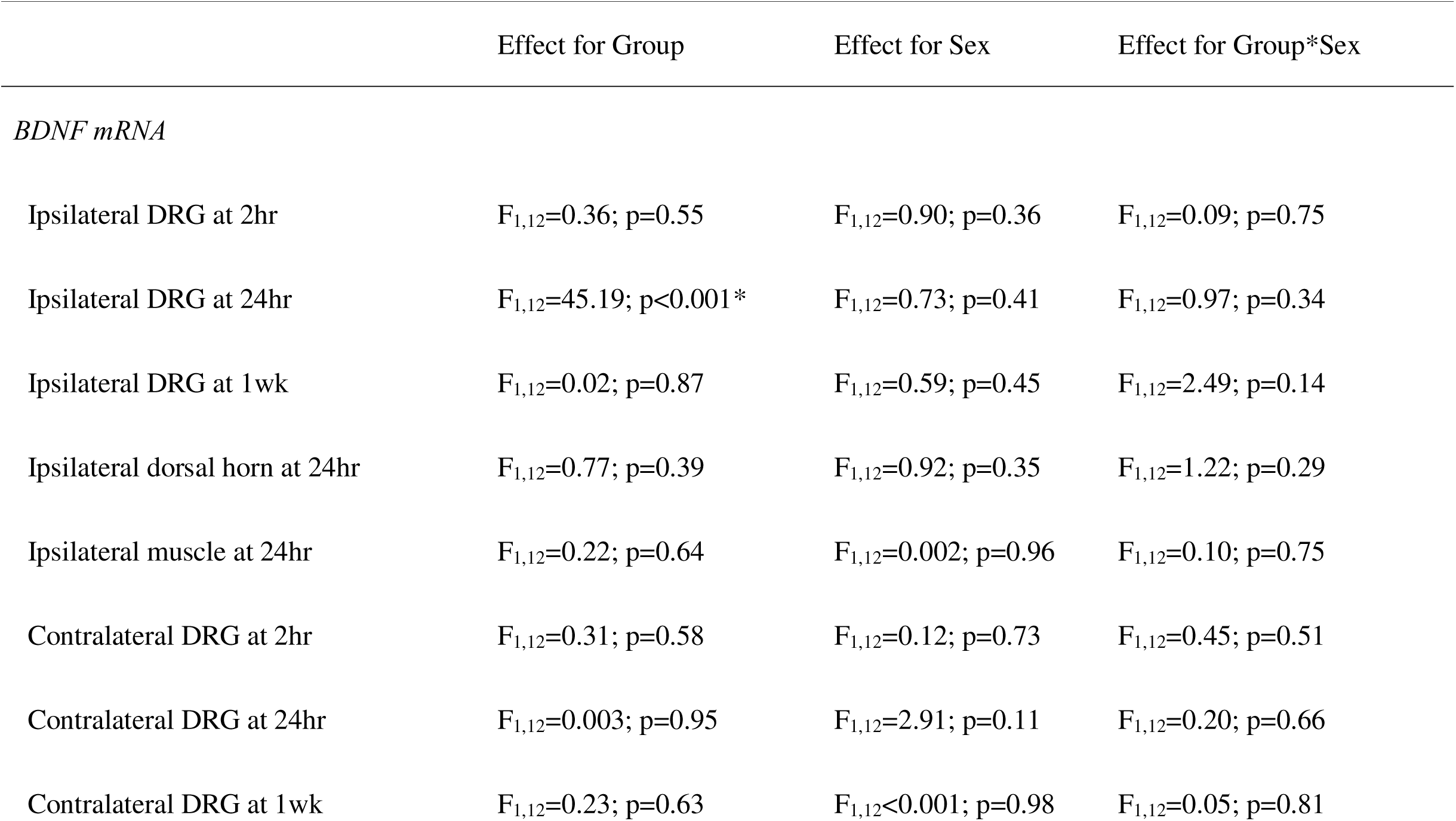
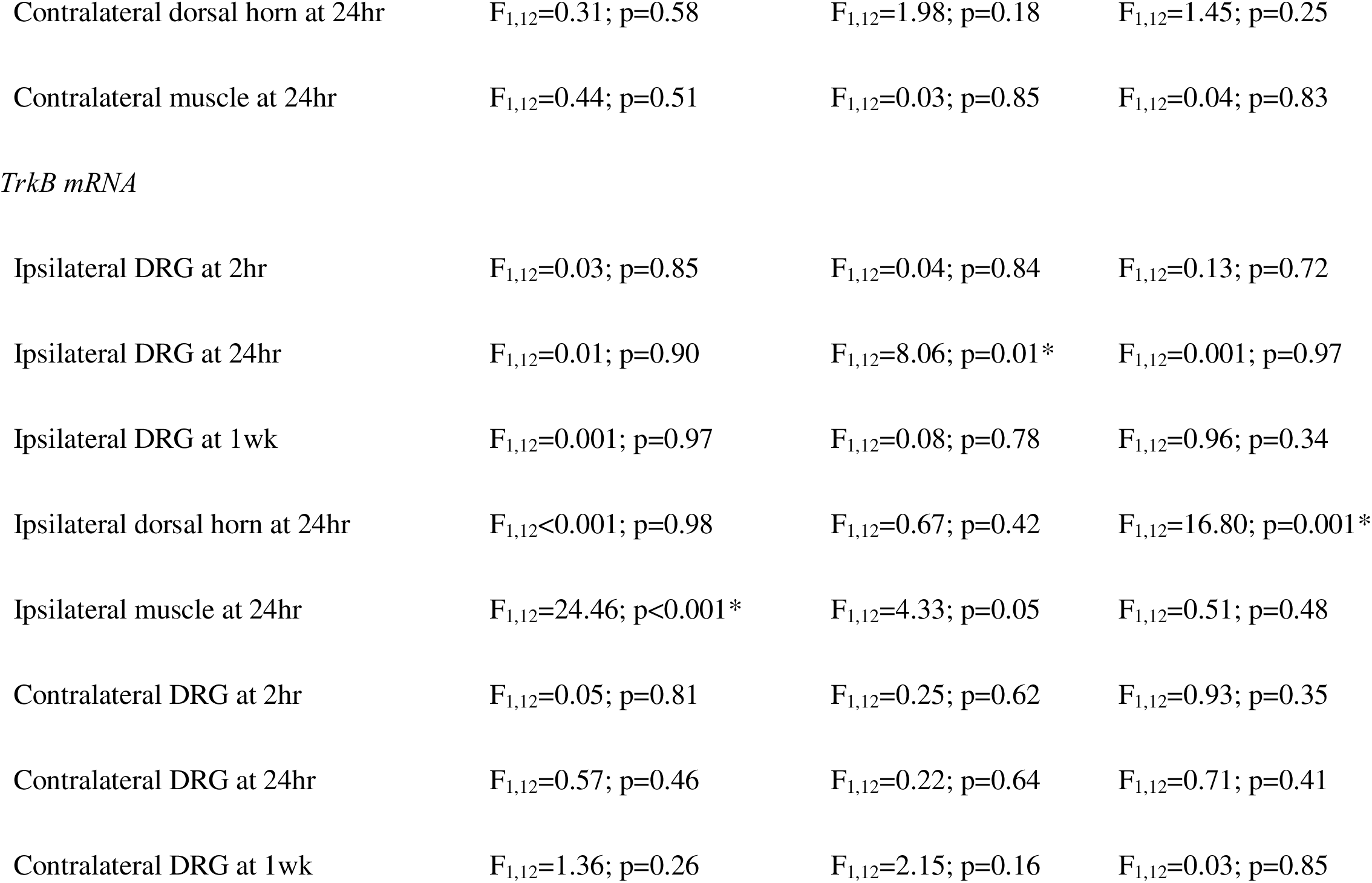
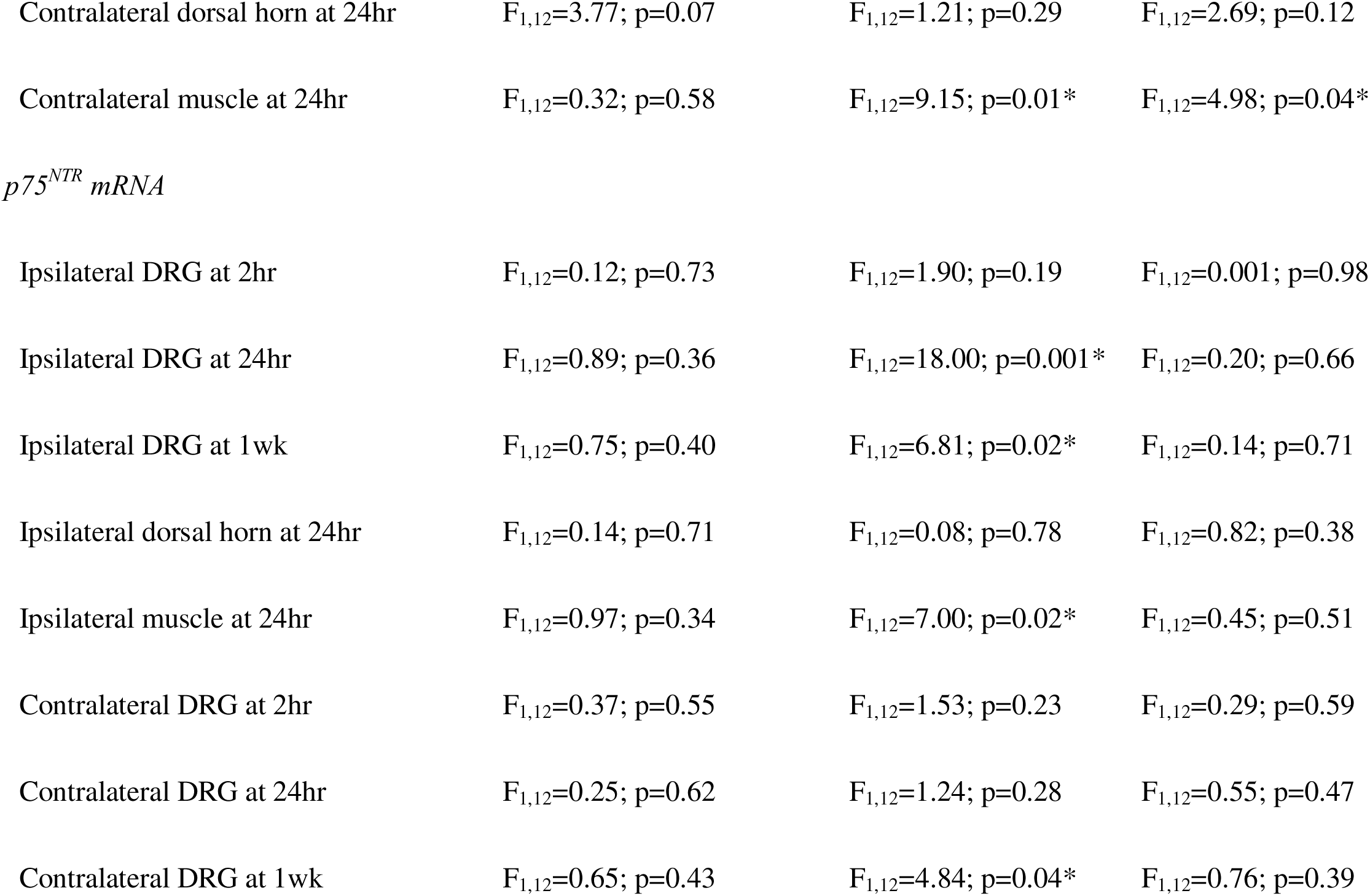
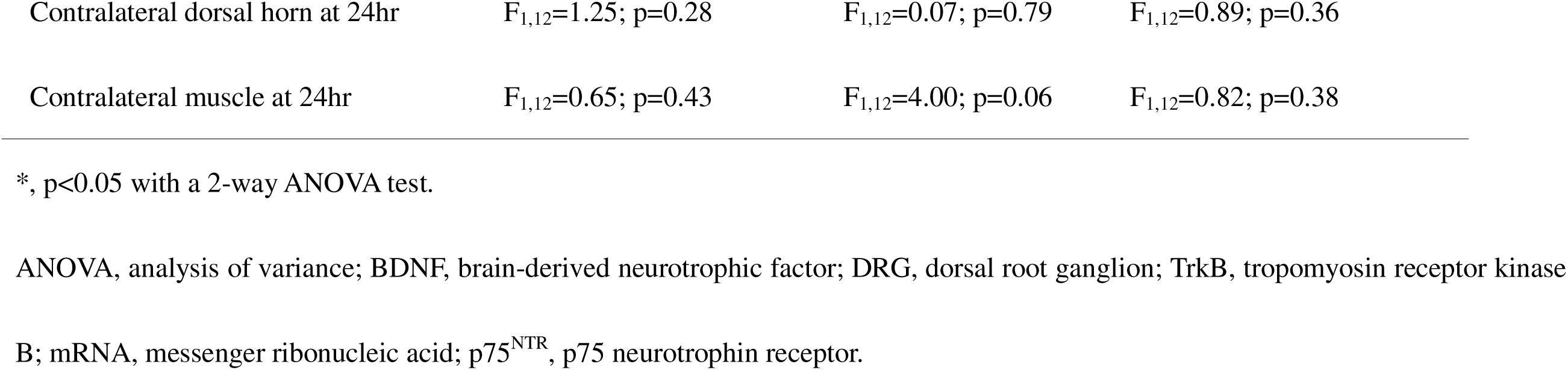
Statistical results (results for a 2-way analysis of variance test for mRNA expression).

|  | Effect for Group | Effect for Sex | Effect for Group*Sex |
| --- | --- | --- | --- |
| *BDNF mRNA* |  |  |  |
| Ipsilateral DRG at 2hr | $F_{1,12}=0.36$; $p=0.55$ | $F_{1,12}=0.90$; $p=0.36$ | $F_{1,12}=0.09$; $p=0.75$ |
| Ipsilateral DRG at 24hr | $F_{1,12}=45.19$; $p<0.001^*$ | $F_{1,12}=0.73$; $p=0.41$ | $F_{1,12}=0.97$; $p=0.34$ |
| Ipsilateral DRG at 1wk | $F_{1,12}=0.02$; $p=0.87$ | $F_{1,12}=0.59$; $p=0.45$ | $F_{1,12}=2.49$; $p=0.14$ |
| Ipsilateral dorsal horn at 24hr | $F_{1,12}=0.77$; $p=0.39$ | $F_{1,12}=0.92$; $p=0.35$ | $F_{1,12}=1.22$; $p=0.29$ |
| Ipsilateral muscle at 24hr | $F_{1,12}=0.22$; $p=0.64$ | $F_{1,12}=0.002$; $p=0.96$ | $F_{1,12}=0.10$; $p=0.75$ |
| Contralateral DRG at 2hr | $F_{1,12}=0.31$; $p=0.58$ | $F_{1,12}=0.12$; $p=0.73$ | $F_{1,12}=0.45$; $p=0.51$ |
| Contralateral DRG at 24hr | $F_{1,12}=0.003$; $p=0.95$ | $F_{1,12}=2.91$; $p=0.11$ | $F_{1,12}=0.20$; $p=0.66$ |
| Contralateral DRG at 1wk | $F_{1,12}=0.23$; $p=0.63$ | $F_{1,12}<0.001$; $p=0.98$ | $F_{1,12}=0.05$; $p=0.81$ |
| Contralateral dorsal horn at 24hr | $F_{1,12}=0.31$; $p=0.58$ | $F_{1,12}=1.98$; $p=0.18$ | $F_{1,12}=1.45$; $p=0.25$ |
| Contralateral muscle at 24hr | $F_{1,12}=0.44$; $p=0.51$ | $F_{1,12}=0.03$; $p=0.85$ | $F_{1,12}=0.04$; $p=0.83$ |
| *TrkB mRNA* |  |  |  |
| Ipsilateral DRG at 2hr | $F_{1,12}=0.03$; $p=0.85$ | $F_{1,12}=0.04$; $p=0.84$ | $F_{1,12}=0.13$; $p=0.72$ |
| Ipsilateral DRG at 24hr | $F_{1,12}=0.01$; $p=0.90$ | $F_{1,12}=8.06$; $p=0.01^*$ | $F_{1,12}=0.001$; $p=0.97$ |
| Ipsilateral DRG at 1wk | $F_{1,12}=0.001$; $p=0.97$ | $F_{1,12}=0.08$; $p=0.78$ | $F_{1,12}=0.96$; $p=0.34$ |
| Ipsilateral dorsal horn at 24hr | $F_{1,12}<0.001$; $p=0.98$ | $F_{1,12}=0.67$; $p=0.42$ | $F_{1,12}=16.80$; $p=0.001^*$ |
| Ipsilateral muscle at 24hr | $F_{1,12}=24.46$; $p<0.001^*$ | $F_{1,12}=4.33$; $p=0.05$ | $F_{1,12}=0.51$; $p=0.48$ |
| Contralateral DRG at 2hr | $F_{1,12}=0.05$; $p=0.81$ | $F_{1,12}=0.25$; $p=0.62$ | $F_{1,12}=0.93$; $p=0.35$ |
| Contralateral DRG at 24hr | $F_{1,12}=0.57$; $p=0.46$ | $F_{1,12}=0.22$; $p=0.64$ | $F_{1,12}=0.71$; $p=0.41$ |
| Contralateral DRG at 1wk | $F_{1,12}=1.36$; $p=0.26$ | $F_{1,12}=2.15$; $p=0.16$ | $F_{1,12}=0.03$; $p=0.85$ |
| Contralateral dorsal horn at 24hr | $F_{1,12}=3.77$; $p=0.07$ | $F_{1,12}=1.21$; $p=0.29$ | $F_{1,12}=2.69$; $p=0.12$ |
| Contralateral muscle at 24hr | $F_{1,12}=0.32$; $p=0.58$ | $F_{1,12}=9.15$; $p=0.01^*$ | $F_{1,12}=4.98$; $p=0.04^*$ |
| *p75<sup>NTR</sup> mRNA* |  |  |  |
| Ipsilateral DRG at 2hr | $F_{1,12}=0.12$; $p=0.73$ | $F_{1,12}=1.90$; $p=0.19$ | $F_{1,12}=0.001$; $p=0.98$ |
| Ipsilateral DRG at 24hr | $F_{1,12}=0.89$; $p=0.36$ | $F_{1,12}=18.00$; $p=0.001^*$ | $F_{1,12}=0.20$; $p=0.66$ |
| Ipsilateral DRG at 1wk | $F_{1,12}=0.75$; $p=0.40$ | $F_{1,12}=6.81$; $p=0.02^*$ | $F_{1,12}=0.14$; $p=0.71$ |
| Ipsilateral dorsal horn at 24hr | $F_{1,12}=0.14$; $p=0.71$ | $F_{1,12}=0.08$; $p=0.78$ | $F_{1,12}=0.82$; $p=0.38$ |
| Ipsilateral muscle at 24hr | $F_{1,12}=0.97$; $p=0.34$ | $F_{1,12}=7.00$; $p=0.02^*$ | $F_{1,12}=0.45$; $p=0.51$ |
| Contralateral DRG at 2hr | $F_{1,12}=0.37$; $p=0.55$ | $F_{1,12}=1.53$; $p=0.23$ | $F_{1,12}=0.29$; $p=0.59$ |
| Contralateral DRG at 24hr | $F_{1,12}=0.25$; $p=0.62$ | $F_{1,12}=1.24$; $p=0.28$ | $F_{1,12}=0.55$; $p=0.47$ |
| Contralateral DRG at 1wk | $F_{1,12}=0.65$; $p=0.43$ | $F_{1,12}=4.84$; $p=0.04^*$ | $F_{1,12}=0.76$; $p=0.39$ |
| Contralateral dorsal horn at 24hr | $F_{1,12}=1.25$; $p=0.28$ | $F_{1,12}=0.07$; $p=0.79$ | $F_{1,12}=0.89$; $p=0.36$ |
| Contralateral muscle at 24hr | $F_{1,12}=0.65$; $p=0.43$ | $F_{1,12}=4.00$; $p=0.06$ | $F_{1,12}=0.82$; $p=0.38$ |

\*, $p<0.05$ with a 2-way ANOVA test.
ANOVA, analysis of variance; BDNF, brain-derived neurotrophic factor; DRG, dorsal root ganglion; TrkB, tropomyosin receptor kinase
B; mRNA, messenger ribonucleic acid; $p75^{\text{NTR}}$ , $p75$ neurotrophin receptor.

For protein, we examined the density of BDNF immunohistochemical staining in the DRG and the spinal cord 24hr after induction of the model, the time point when there was increased mRNA expression and when the animals were responsive to BDNF inhibitors. (**Fig. 4**)(**Table 6**). BDNF staining density 24hr after induction of the activity-induced muscle pain was significantly higher when compared to pain free controls for the ipsilateral L4–L6 DRG (p<0.001*, d=1.44) and the spinal dorsal horn lamina I-II (p=0.003*, d=1.16), but not spinal dorsal horn lamina III-IV. No changes in BDNF density were observed on the contralateral side.

**Fig. 4.**
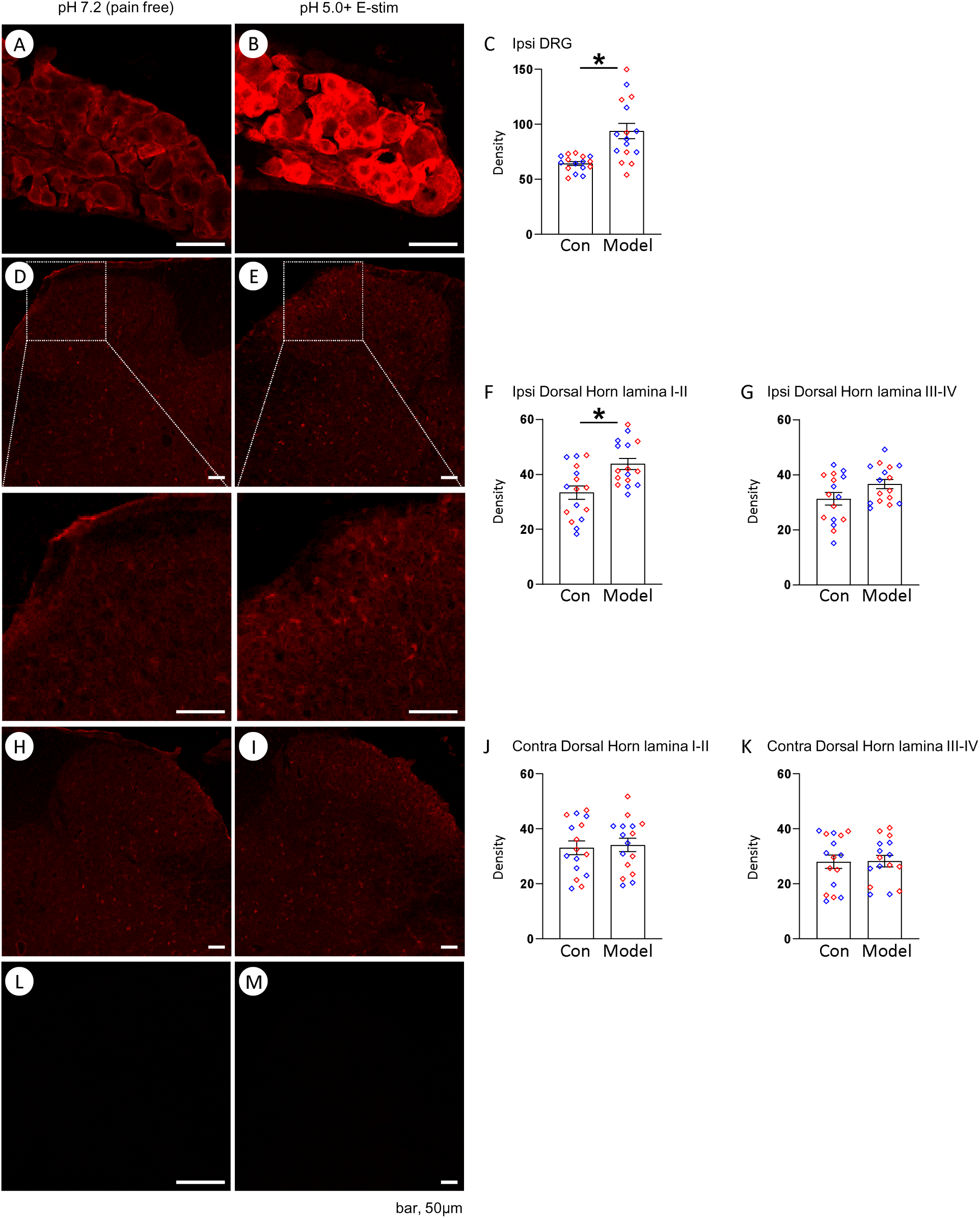
BDNF protein expressions after induction of the activity-induced muscle pain. The BDNF density was significantly increased in ipsilateral L4–L6 DRG (A–C), and spinal dorsal horn lamina I-II (D–G), but not lamina III-IV, 24hr after induction of the model, when compared to pain free controls. No changes were observed in contralateral side (H–K). No antibody control staining for ipsilateral L4–L6 DRG (L), or spinal dorsal horn (M) were shown. Blue plots represent male data, Red plots represent female data. Data is reported as mean ± S.E.M. N=16 in each group. *, p<0.05. BDNF, brain-derived neurotrophic factor; DRG, dorsal root ganglion.

**Table 6.** Statistical results (results for a 2-way analysis of variance test for protein expression).

|  | Effect for Group | Effect for Sex | Effect for Group*Sex |
| --- | --- | --- | --- |
| *BDNF protein density* |  |  |  |
| Ipsilateral DRG at 24hr | $F_{1,28}=15.68$; $p<0.001$* | $F_{1,28}=0.009$; $p=0.92$ | $F_{1,28}=0.05$; $p=0.82$ |
| Ipsilateral dorsal horn lamina I-II at 24hr | $F_{1,28}=10.18$; $p=0.003$* | $F_{1,28}=0.02$; $p=0.87$ | $F_{1,28}=0.15$; $p=0.69$ |
| Ipsilateral dorsal horn lamina III-IV at 24hr | $F_{1,28}=3.45$; $p=0.07$ | $F_{1,28}=0.22$; $p=0.64$ | $F_{1,28}=0.06$; $p=0.80$ |
| Contralateral dorsal horn lamina I-II at 24hr | $F_{1,28}=0.07$; $p=0.79$ | $F_{1,28}=0.25$; $p=0.61$ | $F_{1,28}=0.003$; $p=0.95$ |
| Contralateral dorsal horn lamina III-IV at 24hr | $F_{1,28}=0.005$; $p=0.94$ | $F_{1,28}=0.20$; $p=0.65$ | $F_{1,28}=0.09$; $p=0.76$ |

\*, $p<0.05$ with a 2-way ANOVA test.
ANOVA, analysis of variance; BDNF, brain-derived neurotrophic factor; DRG, dorsal root ganglion.

To determine if the increases in BDNF in the DRG were from primary afferent fibers innervating muscle, we retrogradely labelled muscle afferent DRG neurons with Fast Blue. An average of 75.6 ± 10.6 neurons retrogradely labelled from muscle were counted in L4–L6 DRG (3 sections) in each animal (**Fig. 5**)(**Table 7**). No changes were observed for the overall soma size distribution of BDNF positive cells in Fast Blue labelled neurons after induction of the model when compared to controls (pain model, 388.7 ± 11.6 μm^2^; control, 410.5 ± 25.8 μm^2^; p=0.44). The number of BDNF positive cells was significantly increased, when compared to pain free controls, 24hr after induction of the model for Fast Blue labelled DRG neurons (total, p=0.003*, d=1.58; small-, p=0.007*, d=1.40; medium-, p=0.002*, d=1.67; large-sized, p=0.006*, d=1.45), and non-labelled neurons (total; p<0.001*, d=2.19; small-, p=0.001*, d=1.83; medium-, p<0.001*, d=2.18; large-sized, p=0.03*, d=0.98) in each size category. There were no sex differences in the number of labelled DRG neurons in the activity-induced pain model or pain free control group. For BDNF negative cells in Fast Blue labelled neurons, the overall soma size distribution of neurons showed a shift to small-sized cells in the activity-induced pain model (244.2 ± 11.6 μm^2^), when compared to pain free controls (387.6 ± 16.7 μm^2^)(p<0.001*, d=0.58), however, there were no changes in the total number of cells between groups.

**Fig. 5.**
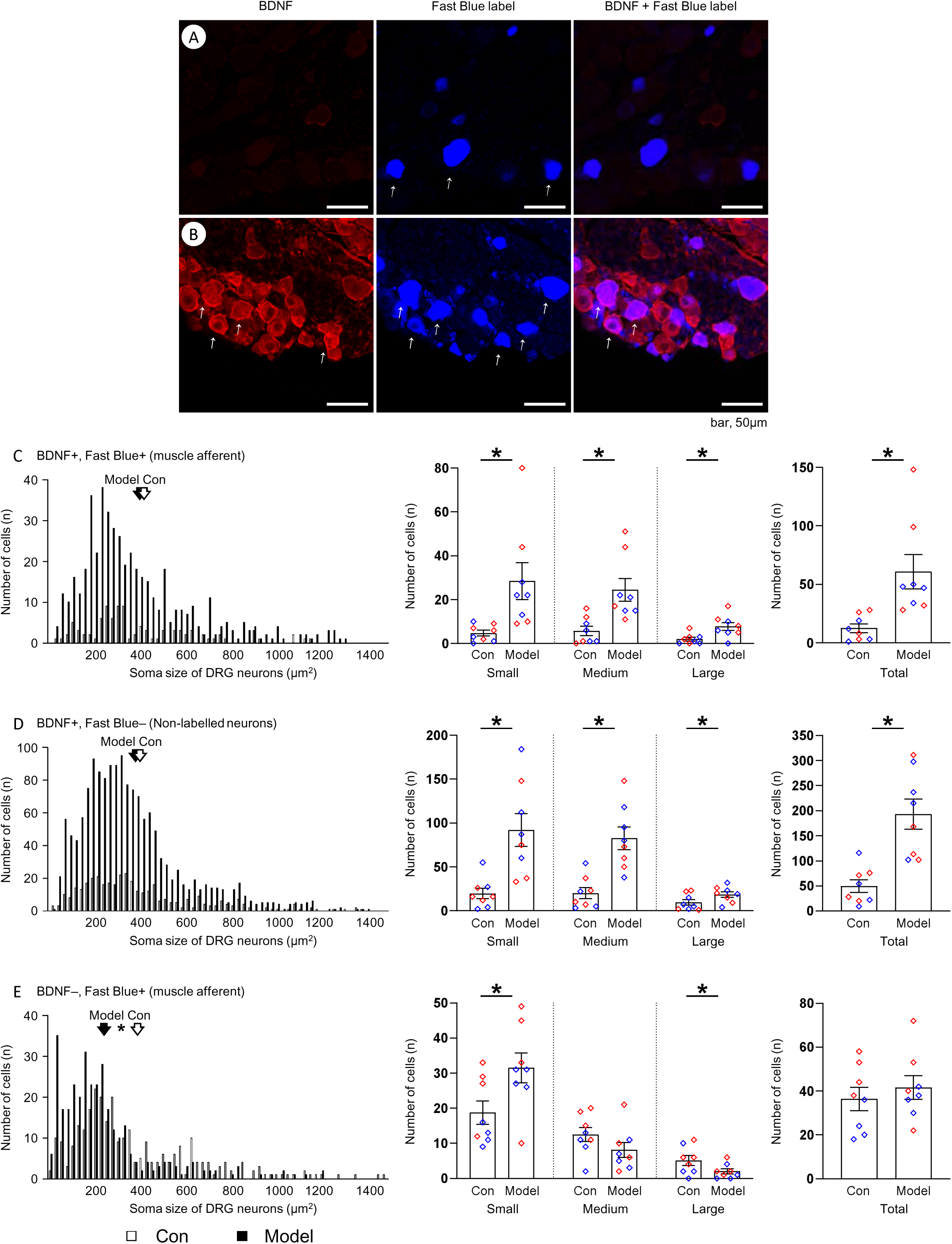
BDNF protein expressions with Fast Blue label after induction of the activity-induced muscle pain. Representative graphs show the BDNF positive or Fast Blue labelled neurons from pain free control (A), or activity-induced pain model (B). The thin white arrows show the positive cells. The number of BDNF positive cells was significantly increased in Fast Blue labelled DRG neuron (C), and Non-labelled neurons (D), 24hr after induction of the activity-induced muscle pain, when compared to pain free controls. The thick white or black arrows show the average soma sizes in each group. No changes were observed in BDNF negative cells in Fast Blue labelled neurons (E). Blue plots represent male data, Red plots represent female data. Data is reported as mean ± S.E.M. N=8 in each group. *, p<0.05. BDNF, brain-derived neurotrophic factor; DRG, dorsal root ganglion.

**Table 7.** Statistical results (results for a 2-way analysis of variance test for protein expression with Fast Blue label).

|  | Effect for Group | Effect for Sex | Effect for Group*Sex |
| --- | --- | --- | --- |
| *Number of DRG neurons (small-sized, <300$\mu\text{m}^2$)* |  |  |  |
| BDNF+, Fast Blue+ neurons (muscle afferent) | $F_{1,12}=7.51$; $p=0.02^*$ | $F_{1,12}=0.96$; $p=0.35$ | $F_{1,12}=0.48$; $p=0.50$ |
| BDNF+, Fast Blue– neurons (non-labelled) | $F_{1,12}=13.20$; $p=0.003^*$ | $F_{1,12}=1.11$; $p=0.31$ | $F_{1,12}=0.69$; $p=0.42$ |
| BDNF–, Fast Blue+ neurons (muscle afferent) | $F_{1,12}=6.38$; $p=0.03^*$ | $F_{1,12}=3.36$; $p=0.09$ | $F_{1,12}=0.55$; $p=0.47$ |
| *Number of DRG neurons (medium-sized, 300–700$\mu\text{m}^2$)* |  |  |  |
| BDNF+, Fast Blue+ neurons (muscle afferent) | $F_{1,12}=11.68$; $p=0.005^*$ | $F_{1,12}=2.00$; $p=0.18$ | $F_{1,12}=0.75$; $p=0.40$ |
| BDNF+, Fast Blue– neurons (non-labelled) | $F_{1,12}=16.26$; $p=0.002^*$ | $F_{1,12}=0.003$; $p=0.96$ | $F_{1,12}=0.003$; $p=0.96$ |
| BDNF–, Fast Blue+ neurons (muscle afferent) | $F_{1,12}=2.59$; $p=0.13$ | $F_{1,12}=3.90$; $p=0.07$ | $F_{1,12}=0.61$; $p=0.45$ |
| *Number of DRG neurons (large-sized, >700$\mu\text{m}^2$)* |  |  |  |
| BDNF+, Fast Blue+ neurons (muscle afferent) | $F_{1,12}=9.83$; $p=0.009^*$ | $F_{1,12}=3.64$; $p=0.08$ | $F_{1,12}=0.67$; $p=0.43$ |
| BDNF+, Fast Blue– neurons (non-labelled) | $F_{1,12}=3.44$; $p=0.09$ | $F_{1,12}=0.08$; $p=0.78$ | $F_{1,12}=0.50$; $p=0.49$ |
| BDNF–, Fast Blue+ neurons (muscle afferent) | $F_{1,12}=4.21$; $p=0.06$ | $F_{1,12}=2.43$; $p=0.15$ | $F_{1,12}=0.33$; $p=0.58$ |
| *Number of DRG neurons (total)* |  |  |  |
| BDNF+, Fast Blue+ neurons (muscle afferent) | $F_{1,12}=10.35$; $p=0.007^*$ | $F_{1,12}=1.73$; $p=0.21$ | $F_{1,12}=0.66$; $p=0.43$ |
| BDNF+, Fast Blue– neurons (non-labelled) | $F_{1,12}=17.35$; $p=0.001^*$ | $F_{1,12}=0.35$; $p=0.56$ | $F_{1,12}=0.30$; $p=0.59$ |
| BDNF–, Fast Blue+ neurons (muscle afferent) | $F_{1,12}=0.71$; $p=0.41$ | $F_{1,12}=7.51$; $p=0.01^*$ | $F_{1,12}=1.18$; $p=0.29$ |

\*, $p<0.05$ with a 2-way ANOVA test.
ANOVA, analysis of variance; BDNF, brain-derived neurotrophic factor; DRG, dorsal root ganglion.

### Fatigue metabolites combined with IL-1β enhance expression of BDNF

We tested if fatigue metabolites alone, IL-1β alone, or fatigue metabolites combined with IL-1β increased BDNF expression in the primary DRG neuron. Significant increases in BDNF mRNA were observed in cultured DRG treated with ATP and pH 6.5 (p=0.04*, d=0.88), IL-1β (p=0.01*, d=1.14), or ATP, pH 6.5 and IL-1β (p=0.004*, d=1.56), when compared to untreated controls (**Fig. 6**)(**Table 9**). There was also a significant increase in BDNF protein with the combination of ATP, pH 6.5 and IL-1β (p=0.02*, d=1.05) when compared to untreated controls. Similarly, TrkB mRNA was increased with the combination of ATP, pH 6.5 and IL-1β when compared to untreated controls (p=0.02*, d=1.13). There were no changes in p75^NTR^ mRNA when compared to controls.

**Fig. 6.**
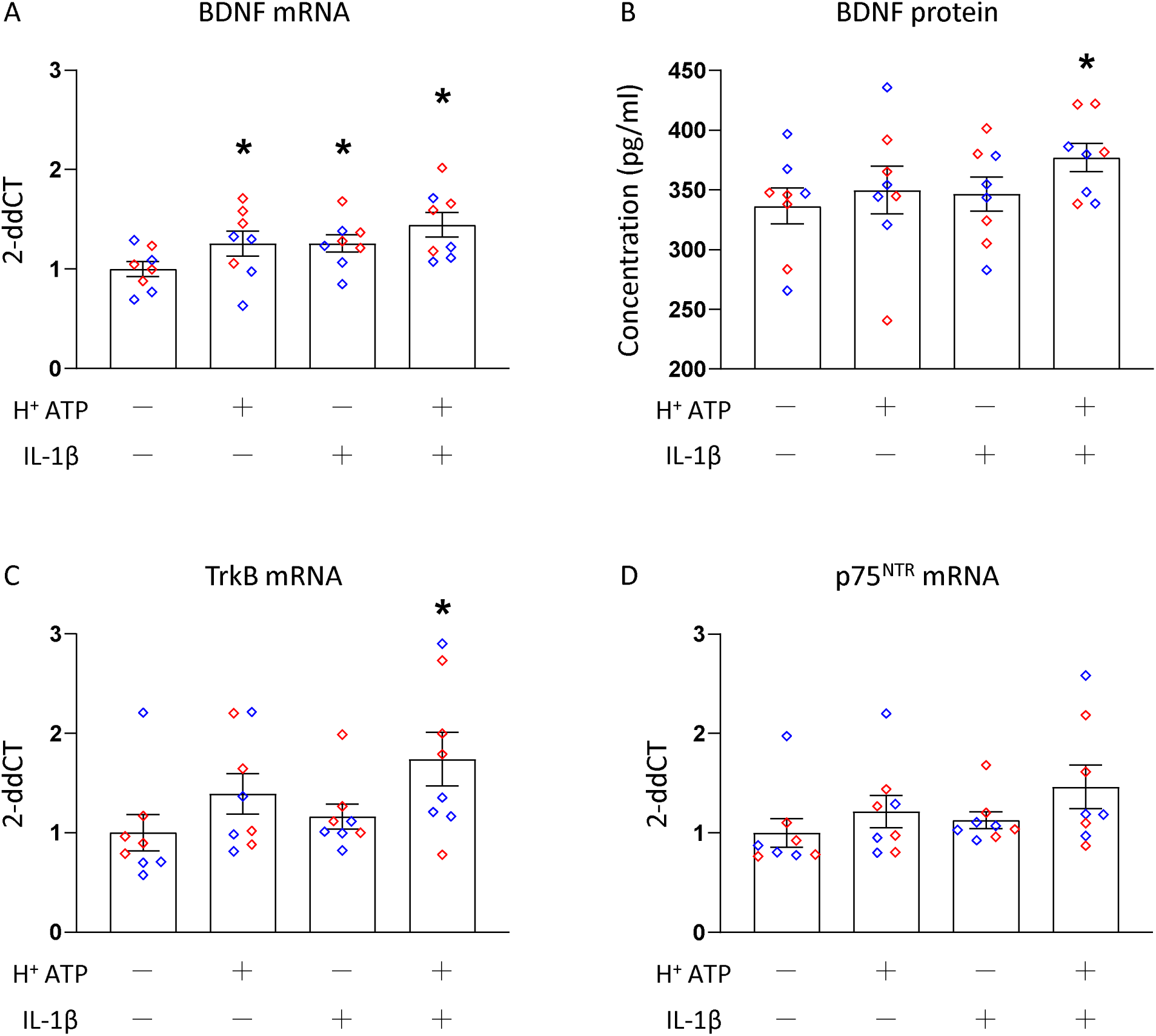
qPCR and ELISA analysis of cultured DRG treated with ATP, pH 6.5, and IL-1β. The cultured DRG with ATP, pH 6.5, and IL-1β showed a significant increase of BDNF mRNA (A) and protein (B), but not TrkB mRNA or p75^NTR^ mRNA (C, D), when compared to the control conditions. Blue plots represent male data, Red plots represent female data. Data is reported as mean ± S.E.M. N=8 in each group. *, p<0.05. ATP, adenosine triphosphate; BDNF, brain-derived neurotrophic factor; DRG, dorsal root ganglion; ELISA, enzyme-linked immunosorbent assay; IL-1β, interleukin-1β; TrkB, tropomyosin receptor kinase B; mRNA, messenger ribonucleic acid; p75^NTR^, p75 neurotrophin receptor; qPCR, quantitative polymerase chain reaction.

**Table 8.** Statistical results (results for a 2-way analysis of variance test for mRNA expression in the animal model of pH4 pain).

|  | Effect for Group | Effect for Sex | Effect for Group*Sex |
| --- | --- | --- | --- |
| *BDNF mRNA* |  |  |  |
| Ipsilateral DRG at 24hr | $F_{1,28}=0.77$; $p=0.38$ | $F_{1,28}=2.55$; $p=0.12$ | $F_{1,28}=0.002$; $p=0.96$ |
| Ipsilateral dorsal horn at 24hr | $F_{1,28}=0.55$; $p=0.46$ | $F_{1,28}=0.04$; $p=0.83$ | $F_{1,28}=0.08$; $p=0.77$ |
| Contralateral DRG at 24hr | $F_{1,28}<0.001$; $p=0.99$ | $F_{1,28}=4.12$; $p=0.05$ | $F_{1,28}=2.39$; $p=0.13$ |
| Contralateral dorsal horn at 24hr | $F_{1,28}=1.74$; $p=0.19$ | $F_{1,28}=2.55$; $p=0.12$ | $F_{1,28}=0.26$; $p=0.60$ |

ANOVA, analysis of variance; BDNF, brain-derived neurotrophic factor; DRG, dorsal root ganglion; mRNA, messenger ribonucleic acid.

**Table 9.** Statistical results (results for a 2-way analysis of variance test for mRNA and protein expression of cultured DRG).

|  | Effect for Group | Effect for Sex | Effect for Group*Sex |
| --- | --- | --- | --- |
| BDNF mRNA | $F_{3,24}=3.68$; $p=0.02^*$ | $F_{1,24}=7.63$; $p=0.01^*$ | $F_{3,24}=0.50$; $p=0.68$ |
| BDNF protein | $F_{3,24}=1.14$; $p=0.35$ | $F_{1,24}=0.002$; $p=0.96$ | $F_{3,24}=0.62$; $p=0.60$ |
| TrkB mRNA | $F_{3,24}=2.23$; $p=0.11$ | $F_{1,24}=0.37$; $p=0.54$ | $F_{3,24}=0.18$; $p=0.90$ |
| p75<sup>NTR</sup> mRNA | $F_{3,24}=1.32$; $p=0.29$ | $F_{1,24}=0.14$; $p=0.71$ | $F_{3,24}=0.29$; $p=0.82$ |

\*, $p<0.05$ with a 2-way ANOVA test.
ANOVA, analysis of variance; BDNF, brain-derived neurotrophic factor; DRG, dorsal root ganglion; TrkB, tropomyosin receptor kinase
B; mRNA, messenger ribonucleic acid; p75<sup>NTR</sup>, p75 neurotrophin receptor.

## Discussion

### Overview

The current data shows that BDNF plays a critical role in development and early phase maintenance of activity-induced muscle pain in male, but not female mice. Specifically, blockade of the BDNF receptor (TrkB) at the site of insult or in the spinal cord reduces development of activity-induced pain in males only. The effects are only observed at the time of induction and 24hr after induction of the model, but not at later time periods, suggesting BDNF plays a role in the early development of activity-induced pain. BDNF is found in primary afferent fibers innervating muscle, and there are increases in mRNA and protein expression after induction of the activity-induced pain model; however, there are no sex differences in expression. In parallel, BDNF expression was increased in the cultured DRG treated with fatigue metabolites, combined with IL-1β, in both male and female mice. Based on these data, we propose BDNF in DRG neurons is both transported to, and released in, the spinal dorsal horn and the muscle to induce activity-induced muscle pain.

### Fatigue metabolites and IL-1β activate DRG to release BDNF and produce hyperalgesia

Fatiguing muscle contractions release ATP and decrease pH [52–54] which can activate purinergic and acid sensing ion channels (ASICs) resulting in hyperalgesia. We previously showed that these fatiguing metabolites activate P2X4 and P2X7 receptors on resident macrophages in the muscle and subsequently increase the release of IL-1β and increase pain [8–10], while blockade of ASIC3 or ASIC1a in muscle prevents activity-induced pain [8, 55]. We have also shown an increase in IL-1β in the gastrocnemius muscle after induction of the activity-induced pain model [10], while blockade of IL-1β prevents the hyperalgesia [9, 10]. Sensory neurons also respond to decreases in pH and ATP through activation of ASICs and P2X receptors on nociceptors [12]. Acidic conditions, combined with ATP, sensitize Group III and IV muscle afferents and DRG neurons [43, 56, 57], and prior studies have shown that combining lactate, ATP and acidic saline produces hyperalgesia in uninjured animals [58]. Using the ischemic perfusion model, prior studies show upregulation of IL-1β/IL1 receptor signaling [59], and this upregulation is dependent on activation of ASIC3 in afferent fibers [60]. Further, injection of IL-1β into the masseter muscle increases expression of P2X3 receptors in DRG, and blockade of P2X3 inhibits hyperalgesia associated with muscle contraction of IL-1β muscle injections [43, 59, 61]. Consistent with these data, the current study demonstrates enhanced expression of BDNF in response to fatigue metabolites and IL-1β and suggests that BDNF may be a downstream mediator of the effects of fatigue metabolites and IL-1β, which can in turn contribute to hyperalgesia.

In parallel, fatigue metabolites and IL-1β were sufficient to increase mRNA expression of BDNF, and their combination resulted in an increase in BDNF protein expression in cultured DRG. A prior study showed that DRG treated with the pro-inflammatory cytokine tumor necrosis factor α, increased BDNF content and release from cultured DRG [44]. The acidic saline model directly activates ASIC3 on muscle afferents to produce pain [62, 63]. Acidic pH 6.0 alone (a similar pH observed in the acid-saline model in muscle after pH 4 injection), but not pH 6.5, induces release of IL-1β from cultured peritoneal macrophages [9, 10, 63]. In the current study we did not show changes in BDNF expression in the acidic saline model suggesting that BDNF expression and release requires activation of multiple receptors. IL-1β could act directly on nociceptors via the type I IL-1 receptor on sensory neurons, or indirectly through release and/or activation of other nociceptive molecules, to produce hyperalgesia [11, 64]. Our prior work also suggests pharmacological blockade of ASIC3 in the muscle prevents development of hyperalgesia, while depletion of ASIC3 from primary afferents has no effect on the activity-induced pain model [8]. Our prior studies, in combination with the current study, suggest that the treatment of cells with fatigue metabolites releases IL-1β protein from macrophages, and then the combination of fatigue metabolites and IL-1β subsequently release BDNF from primary DRG neurons [9, 10, 65].

### BDNF activates a sex-specific mechanism to produce muscle pain

Prior literature on the sex-specific effects of BDNF is controversial and is dependent on the model tested and the site of release of BDNF [25, 27, 30, 31, 39, 66–70]. When injected into peripheral tissues, BDNF produced hyperalgesia [27, 28]; however this was only tested in male mice. Conversely, when injected intrathecally, BDNF produced pain in both sexes [25]. **Table 10** summarizes the data on sex-differences in BDNF in a variety of pain models. Similar to the current study, blockade or genetic deletion of BDNF in males but not females prevented hyperalgesia in models of neuropathic pain and hyperalgesic priming [30, 39, 67, 69]. In contrast, there was a female specific effect for BDNF in visceral pain [66]. In general, the majority of studies demonstrated that the effects of blockade of BDNF were equivalent in both sexes, including models of visceral pain, formalin, and neuropathic pain [25, 27, 31, 39, 68, 69].

**Table 10.**
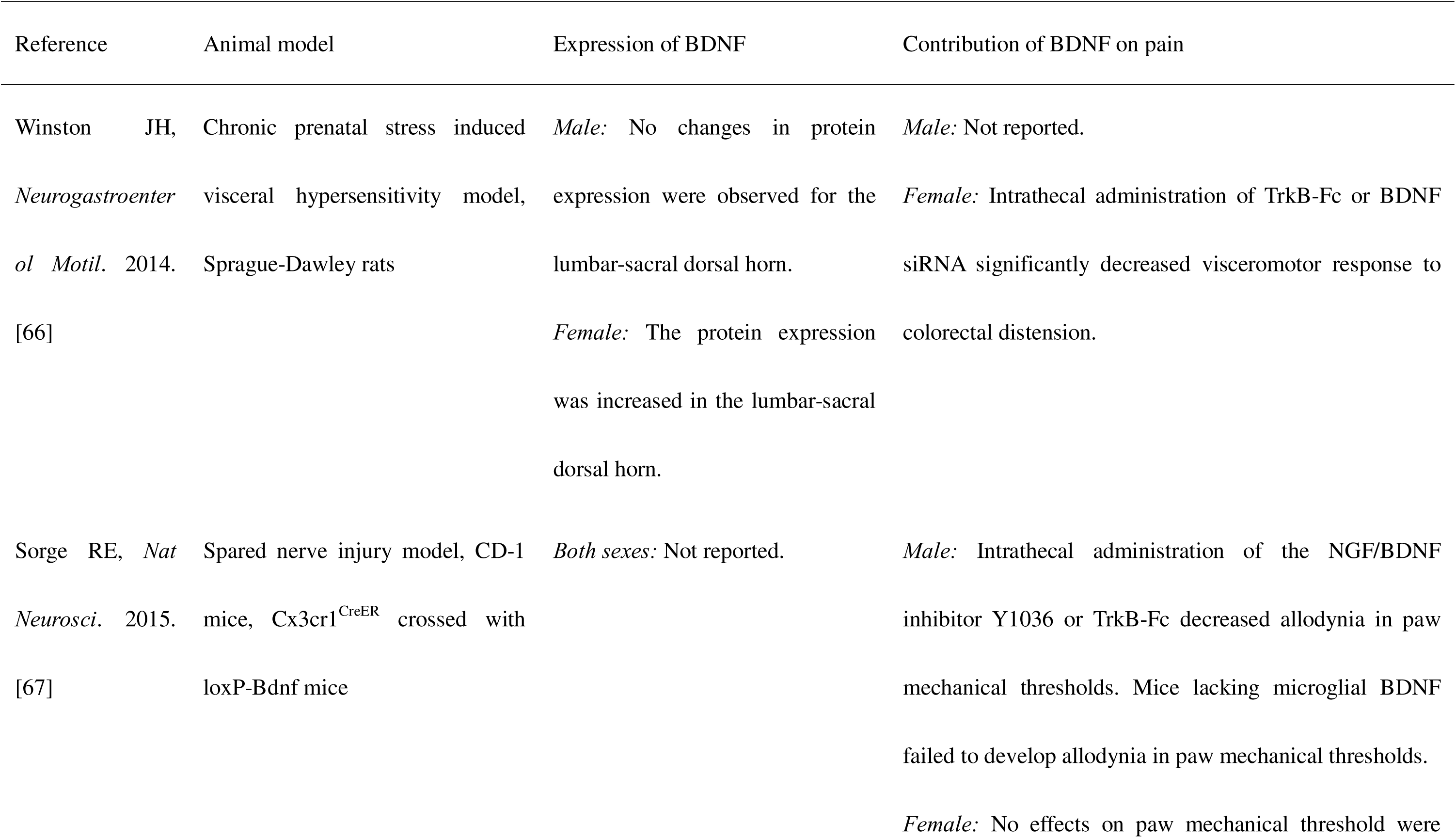
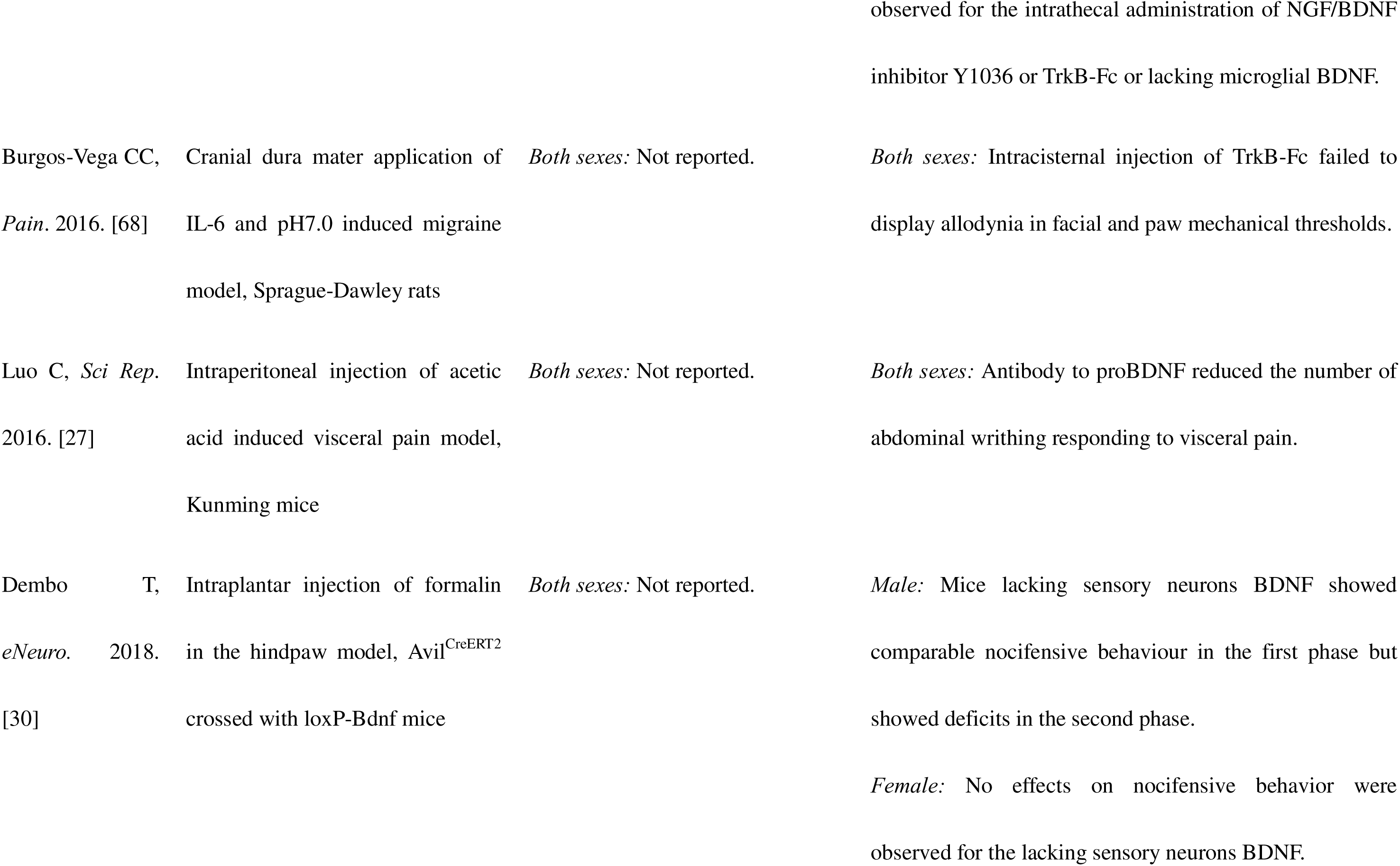
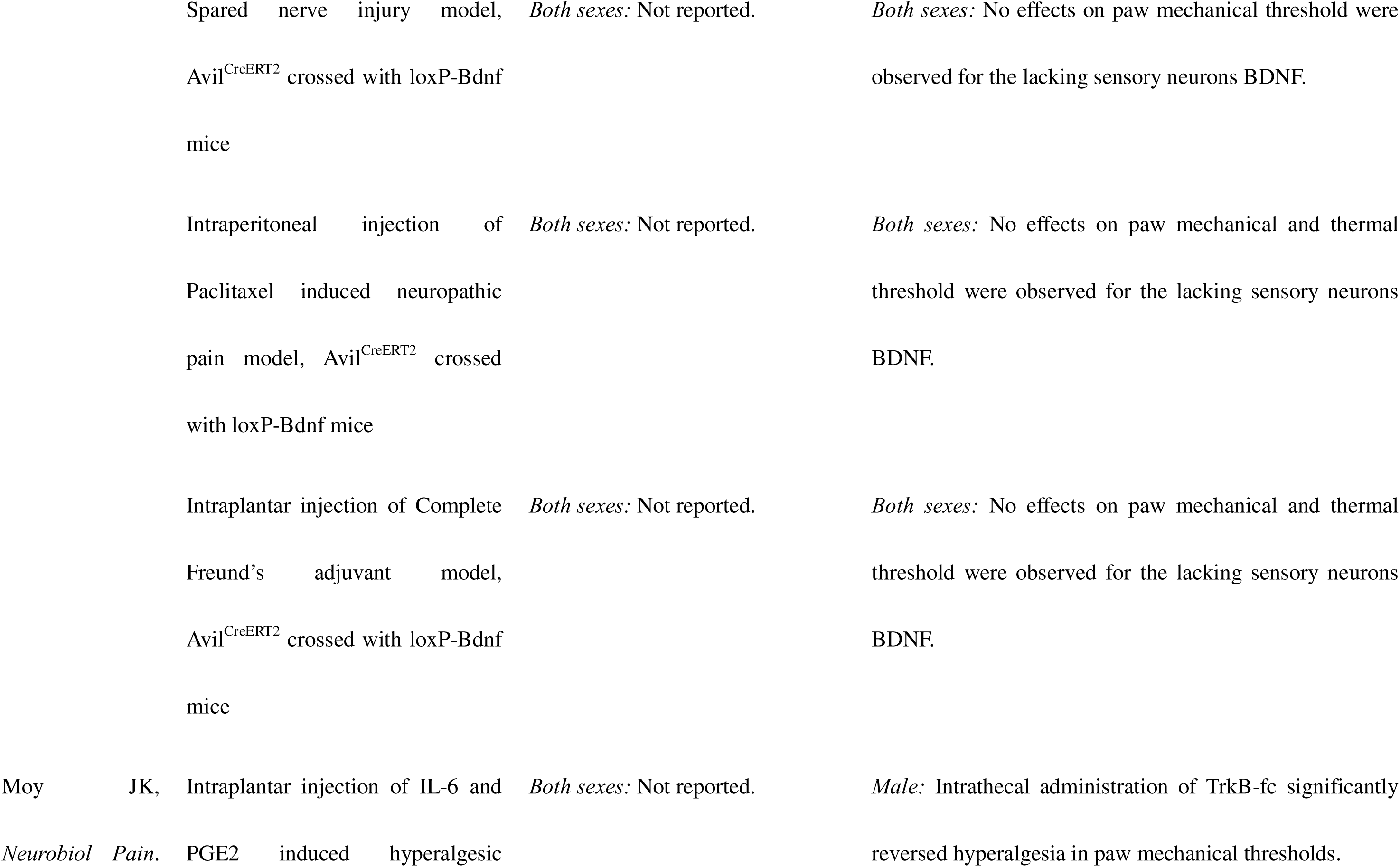
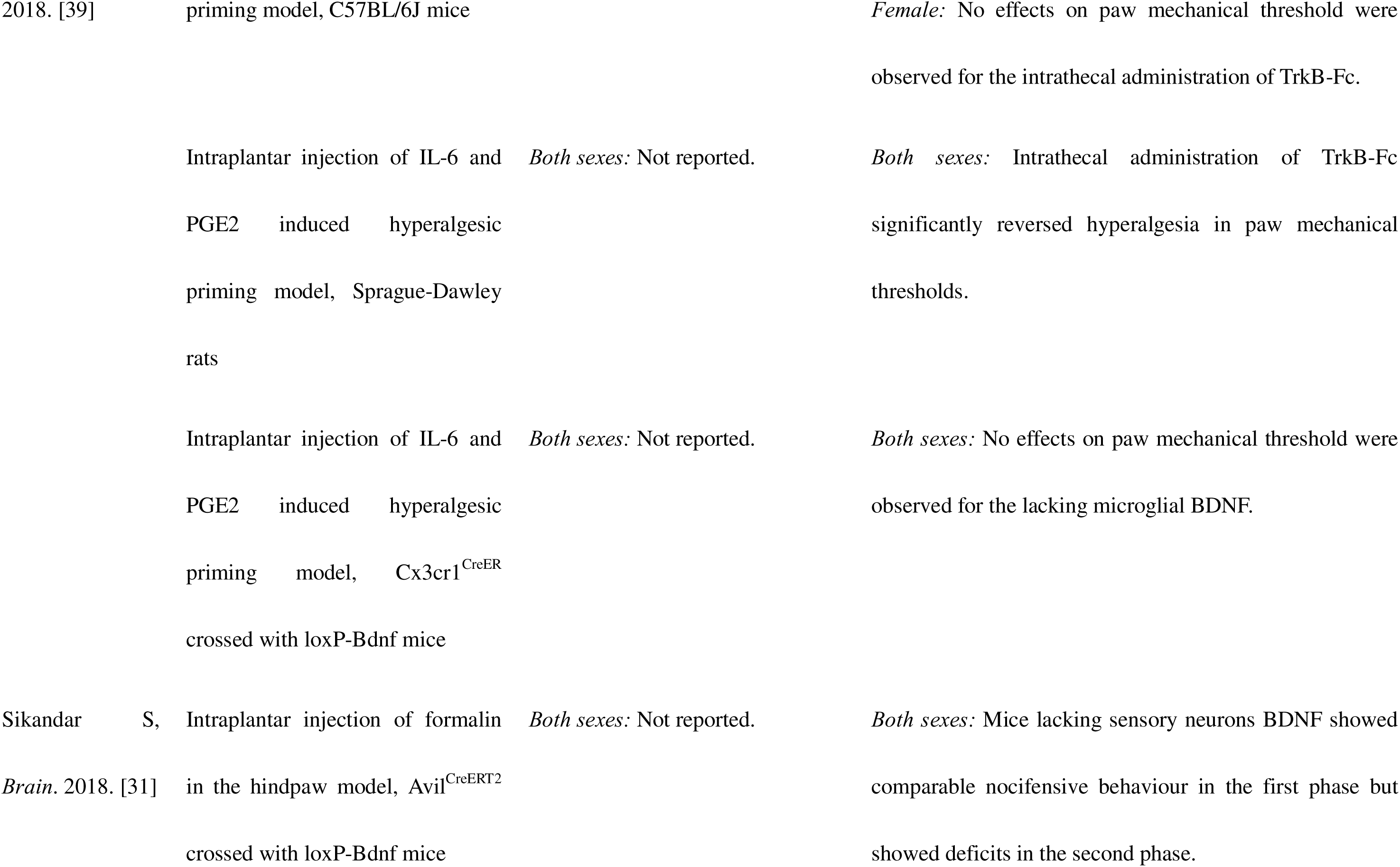
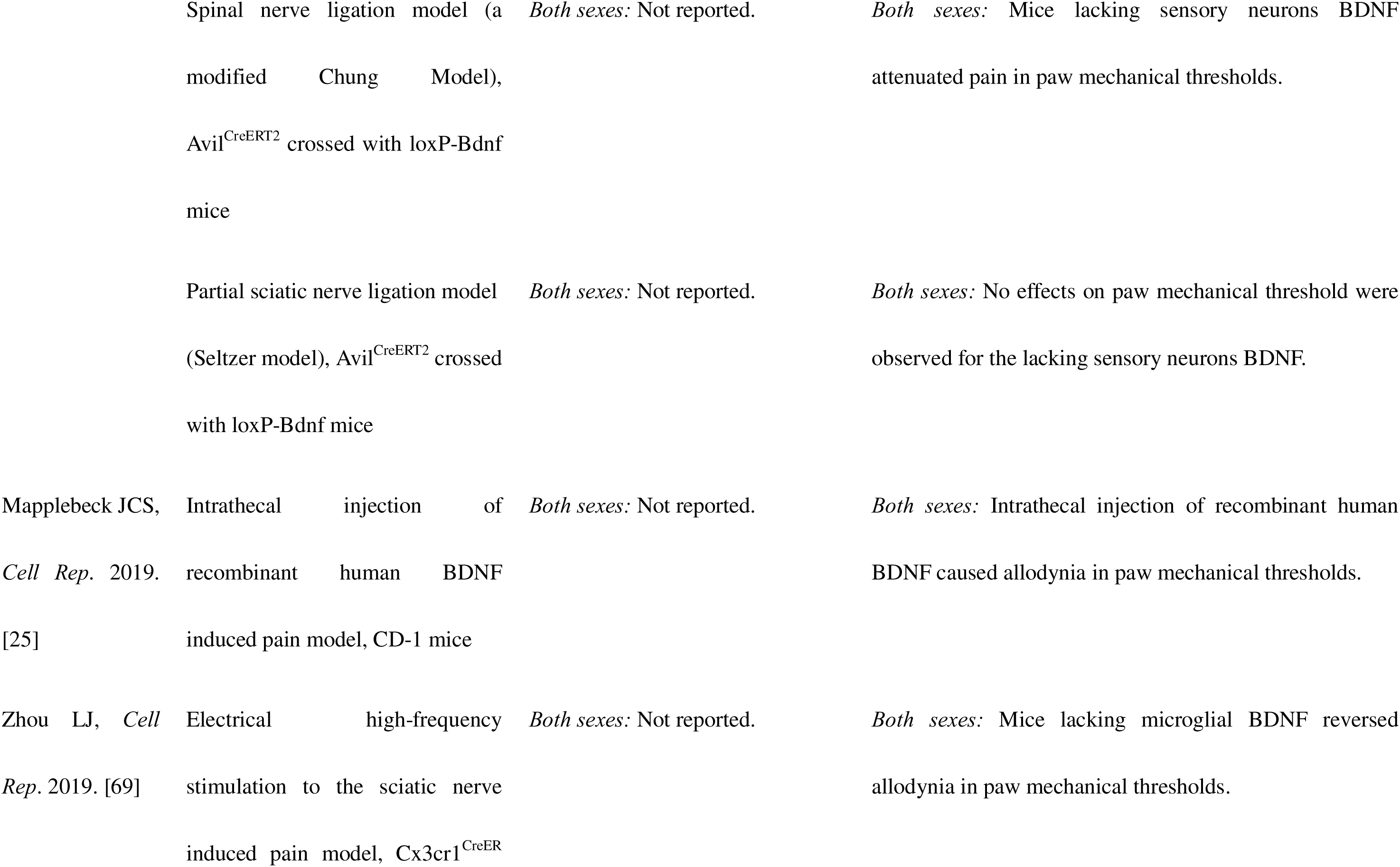
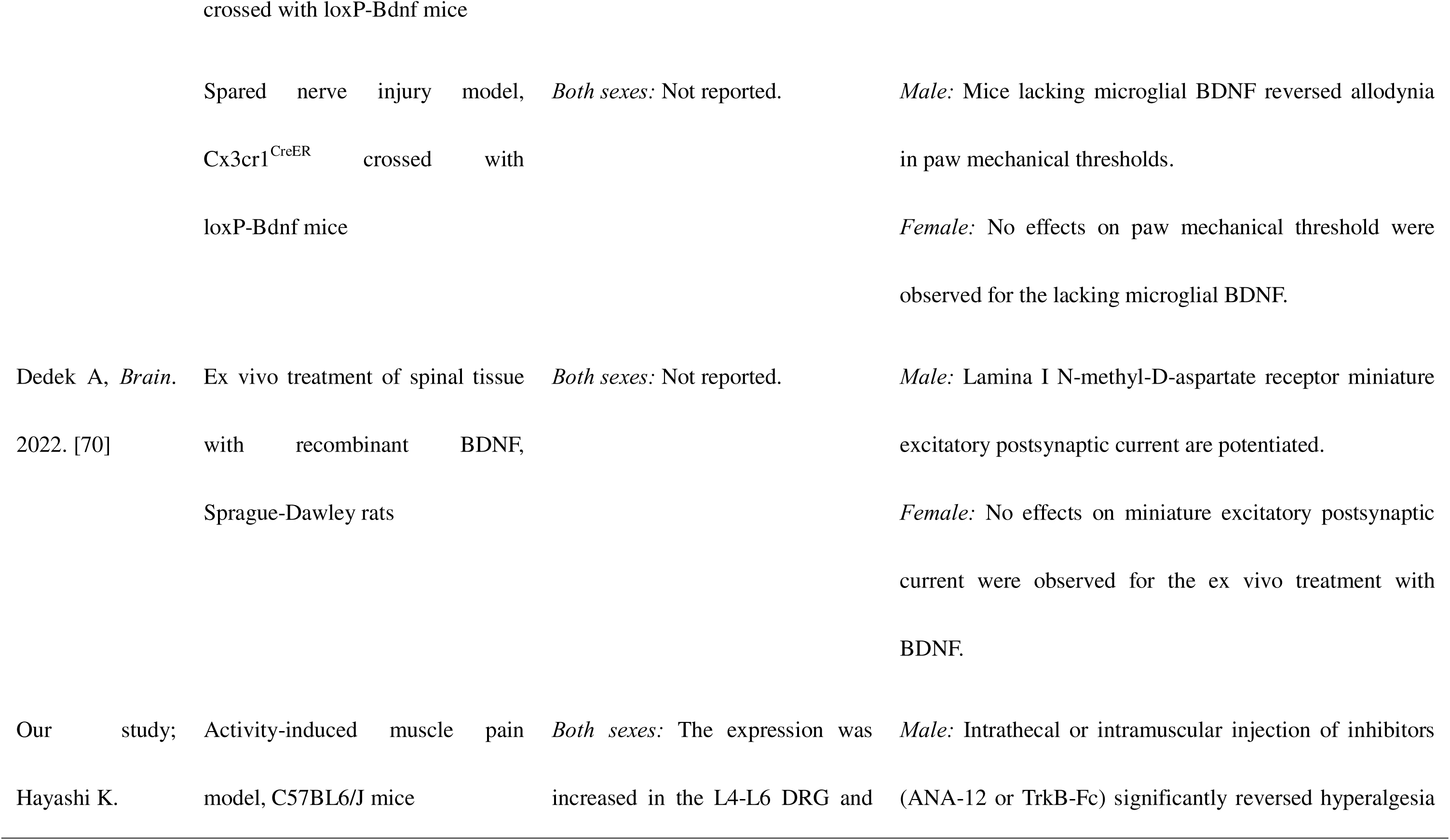
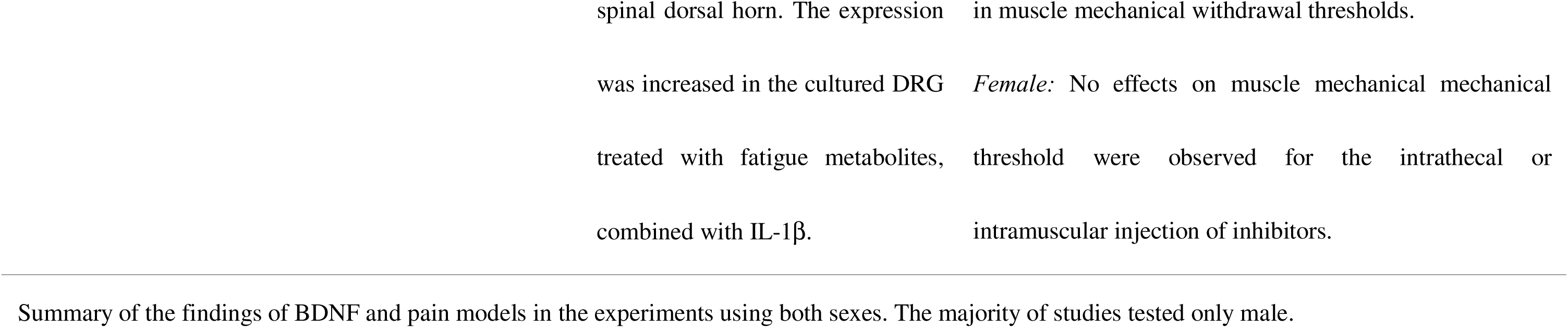
Current findings of sex difference of BDNF and pain models.

BDNF in spinal dorsal horn can be produced by primary afferent fibers or microglia [23, 71, 72]. Genetic studies have started to examine the contributions of primary afferent fibers and microglia on BDNF production and impact. Male, but not female mice, lacking microglial BDNF (Cx3cr1^CreER^ crossed with loxP-Bdnf mice) do not develop neuropathic pain behaviors [67, 69]. Mixed results are found with hyperalgesic priming models and depend on the priming and induction stimuli. For example: 1) removal from sensory neurons prevented carrageenan-PGE2 hyperalgesic priming; 2) BDNF-sequestration at site of injury had no effect on plantar incision-PGE2 hyperalgesic priming; 3) BDNF-sequestration prevented IL-6-PGE2 hyperalgesic priming [31, 39]. Thus, the role of BDNF likely depends on site of action for BDNF (peripheral vs central), targeted cell type (sensory neuron or microglia), sex (male vs female), and model used [25, 27, 29–31, 39, 67–69]. BDNF derived from microglial cells is not likely to contribute to activity-induced pain, because there are no changes in BDNF mRNA in the spinal dorsal horn in either sex. In contrast, the current study supports a role for primary afferent derived BDNF since there is increased release from DRG neurons in response to combining fatigue metabolites with IL-1β, there is upregulation of BDNF in muscle afferents, and peripheral blockade of BDNF receptors reduces hyperalgesia.

The current study suggests that BDNF from muscle afferent DRG neurons, specifically C- or Aδ-fibers, contributes to activity-induced muscle pain in male but not female mice. The soma size distribution of muscle afferent neurons shows no change in BDNF positive cells, although demonstrates a shift to small-sized cells in BDNF negative cells. In contrast to the current study, a size shift to a larger size of BDNF positive cells has been observed in inflammatory and neuropathic pain models, which may contribute to allodynia [18, 73–75]. BDNF, synthesized in the primary DRG neurons, may act as an autocrine or paracrine signal to activate the pre-synaptic TrkB, or p75^NTR^, receptor and regulate synaptic excitability in pain transmission [44]. Consistent with this, the current study shows increases in BDNF not only in muscle DRG neurons but also in non-labelled DRG neurons. In addition, BDNF is anterogradely transported to the spinal dorsal horn to modulate post synaptic responses [13, 17, 18, 20–22]; these effects may be sex-specific. For example, in male, but not female mice, application of BDNF increases expression of glutamate receptor GluN2B, decreases expression of the neuron specific chloride extruder KCC2, and potentiates NMDAR excitatory postsynaptic potentials [70]. It should be noted that BDNF receptors are expressed on myocytes and local immune cells like macrophages can also express BDNF and its receptors [23, 71, 72]. It is therefore also possible that peripheral BDNF is derived from local immune or muscle cells in addition to sensory afferents.

### Sex-specific differences in the activity-induced muscle pain

There are significant sex differences in the generation of pain, however, previous mouse and rat studies have predominately used males only [76]. Gonadectomized and gonadal hormone replacement alter the response of BDNF and the activity-induced pain model [67, 77]. Interestingly, ovariectomy recapitulated the effects of BDNF in female mice normally observed in male mice, showing increased expression of glutamate receptor GluN2B, decreased expression of the neuron specific chloride extruder KCC2, and potentiated NMDAR excitatory postsynaptic potentials [70], suggesting that hormones may modulate the effects of BDNF. Consistent with the role of hormones in modulating pain, we have shown that testosterone protects against the development of widespread pain in the activity-induced pain model [78], with orchiectomy producing widespread pain similar to females and testosterone producing unilateral pain similar to males. Further, we have shown female-specific mechanisms of pain in the activity-induced pain model where there was activation of the antigen processing and presentation pathway in the muscle, and blockade of MHC II attenuated development of muscle hyperalgesia [79]. Centrally, females but not males, also have greater serotonin transporter immunoreactivity in the nucleus raphe magnus [78]. In the muscle, males show a P2X7-dependent hyperalgesia in the activity induced pain model [10]. In contrast, ASIC3 and P2X4 mediate hyperalgesia in the activity-induced pain model in both sexes. Thus, there is a complicated network of pathways that mediate hyperalgesia that may be dependent on the specific molecular pathway involved in the underlying mechanism and the pain model studied.

## Conclusions

A schematic representation of the contribution of BDNF in the activity-induced pain model in male and female mice is shown in **Fig. 7**. The present study uniquely showed there was a sex difference in the contribution of BDNF on pain, while there were no sex differences in the expression changes observed in BDNF. Similarly, we previously showed the same for activation of the P2X7-NLRP3-Caspase-1 pathway where the hyperalgesia depended on activation of the pathway in males, but there were changes in expression of these proteins in both male and females [10]. We propose that there are both shared and unique pathways in male and female mice that contribute to the development of hyperalgesia.

**Fig. 7.**
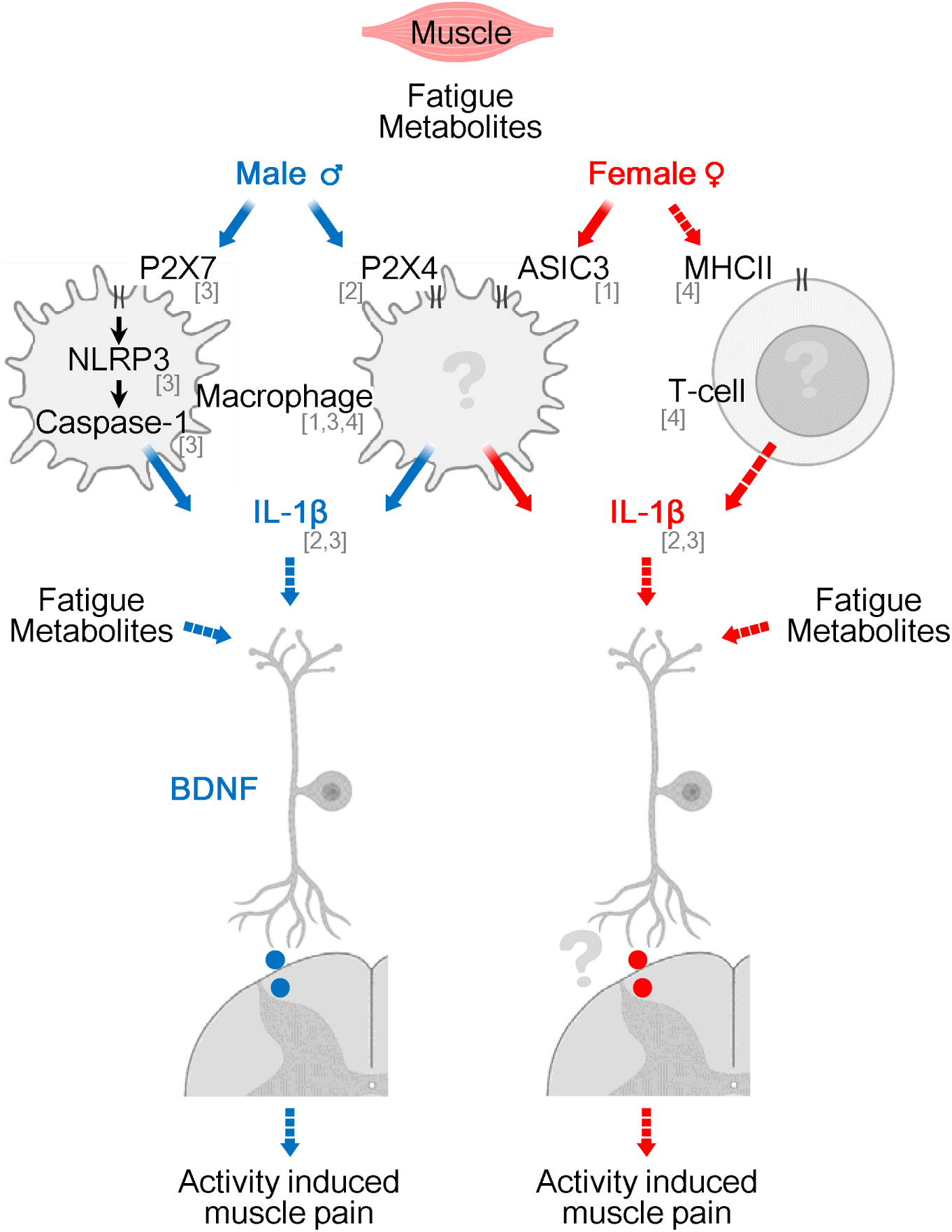
Schematic representation of the contribution of BDNF to the activity-induced pain. The activity-induced muscle pain in early maintenance phases is contributed by BDNF on DRG in male mice. [1] Gregory N, et al. Mol Neurobiol. 2016; [2] Oliveira-Fusaro MC, et al. Mol Neurobiol. 2020; [3] Hayashi K, et al. Pain. 2023; [4] Lesnak JB, et al. Brain Behav Immun. 2023.

## Acknowledgements

This study is supported by the National Institutes of Health AR073187. Dr. Hayashi is supported by Japan Society for the Promotion of Science. The funders played no role in the design, conduct, or reporting of this study. There was no additional external funding received for this study. The authors would like to acknowledge the BioRender.com to create scientific figures.

## Figure Legends

**Supplemental Fig. 1.**
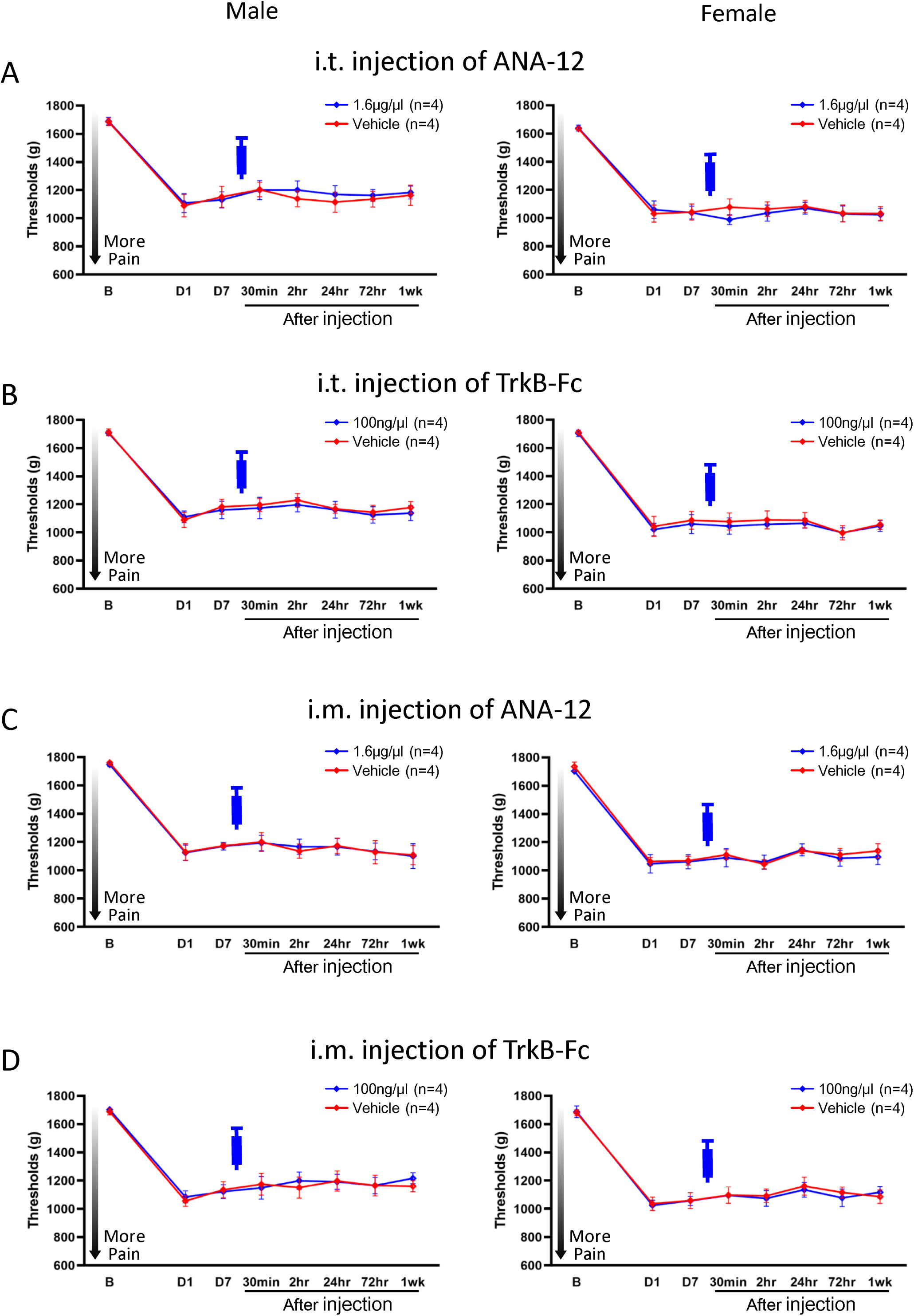
Ipsilateral muscle withdrawal thresholds with pharmacological inhibition intrathecally or intramuscularly 1wk after induction of the model. BDNF inhibitors intrathecally or intramuscularly 1wk after induction of the model had no effect on muscle withdrawal thresholds on ipsilateral side in male or female mice (A–D). Data is reported as mean ± S.E.M. *, versus vehicle control; p<0.05. i.m., intramuscularly; i.t., intrathecally.

**Supplemental Fig. 2.**
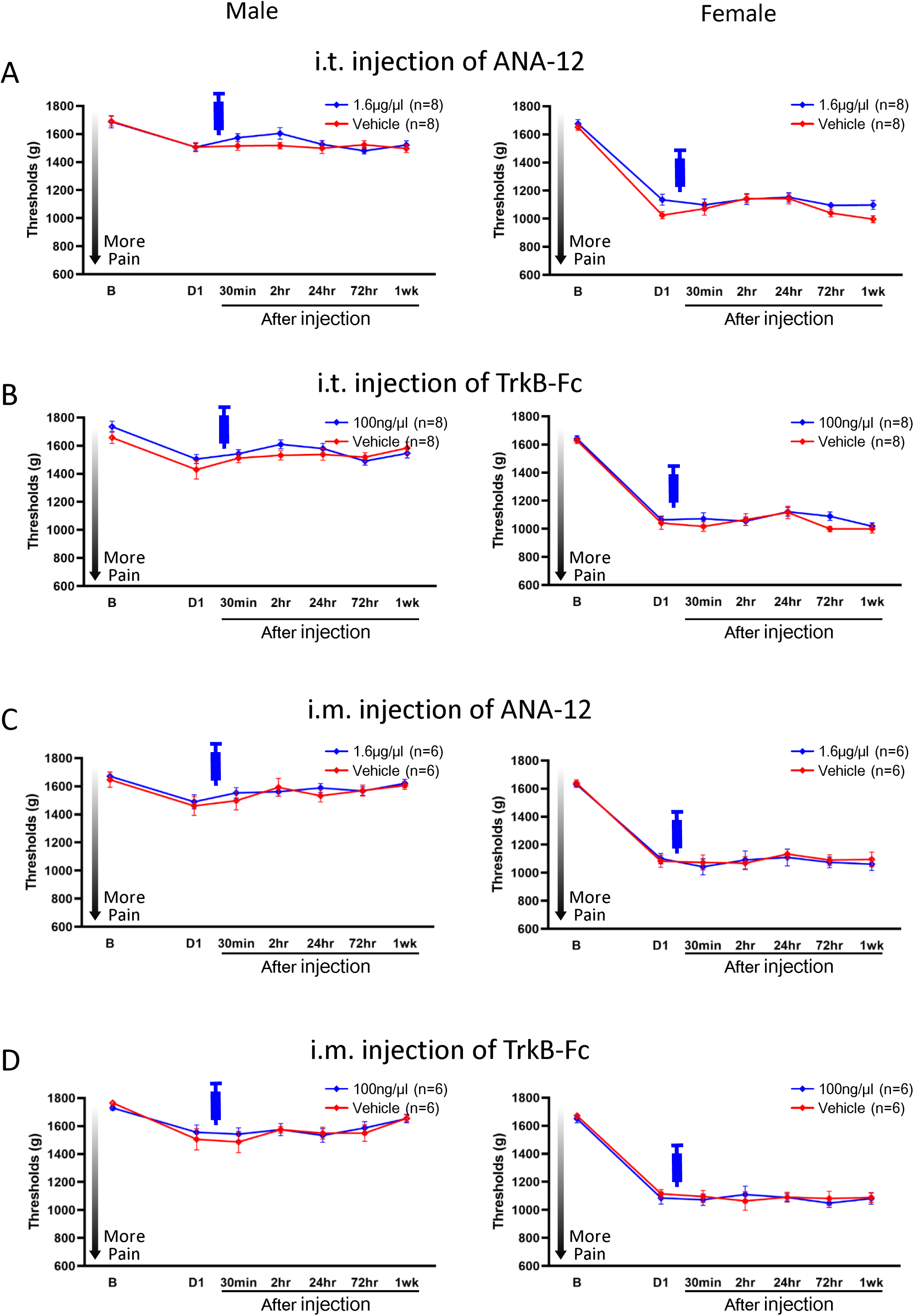
Contralateral muscle withdrawal thresholds with pharmacological inhibition intrathecally or intramuscularly 24hr after induction of the model. BDNF inhibitors intrathecally or intramuscularly 24hr after induction of the model had no effect on muscle withdrawal thresholds on contralateral side in male or female mice (A–D). Data is reported as mean ± S.E.M. *, versus vehicle control; p<0.05. i.m., intramuscularly; i.t., intrathecally.

**Supplemental Fig. 3.**
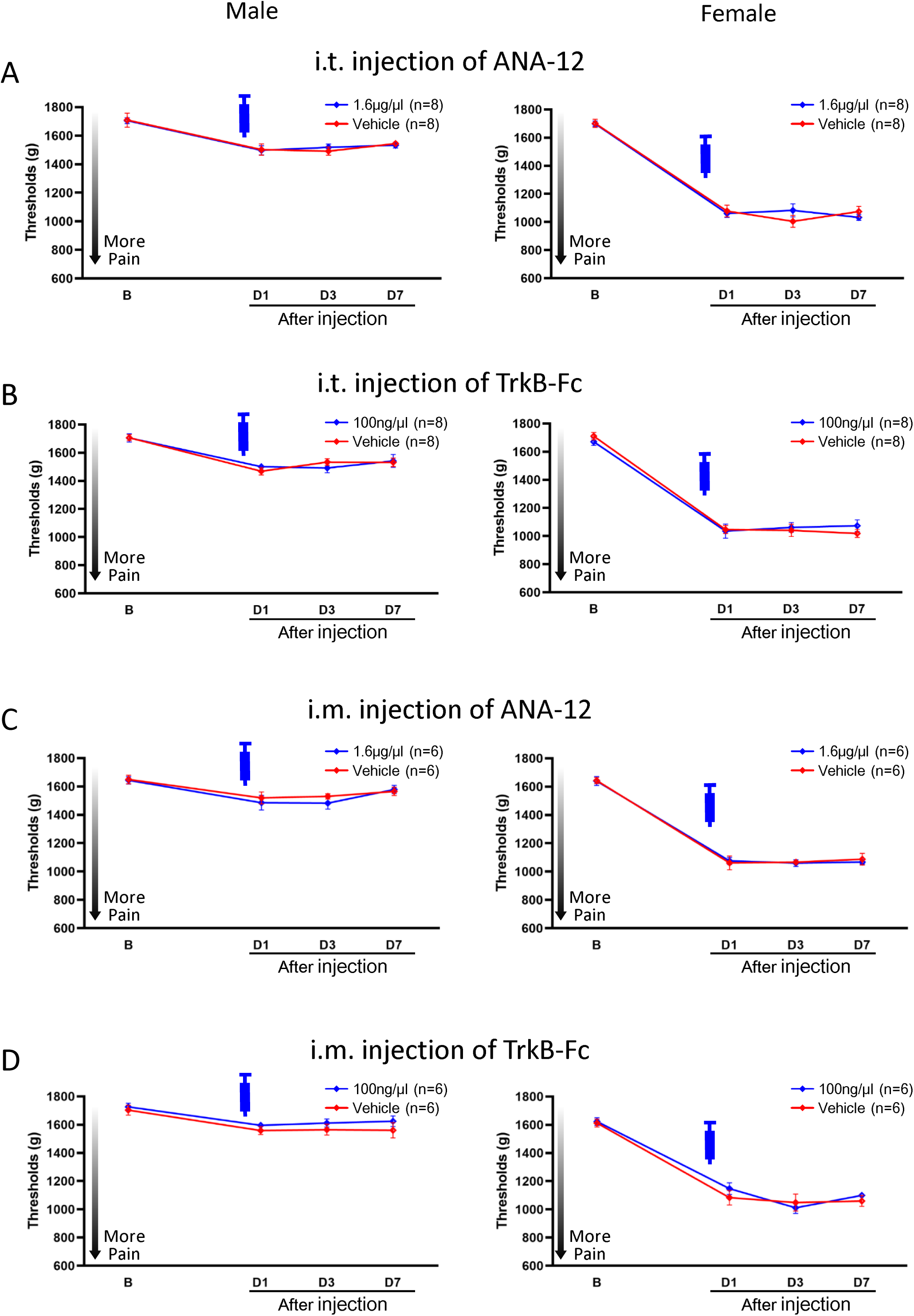
Contralateral muscle withdrawal thresholds with pharmacological inhibition intrathecally or intramuscularly prior to induction of the model. BDNF inhibitors intrathecally or intramuscularly prior to induction of the model had no effect on muscle withdrawal thresholds on contralateral side in male or female mice (A–D). Data is reported as mean ± S.E.M. *, versus vehicle control; p<0.05. i.m., intramuscularly; i.t., intrathecally.

**Supplemental Fig. 4.**
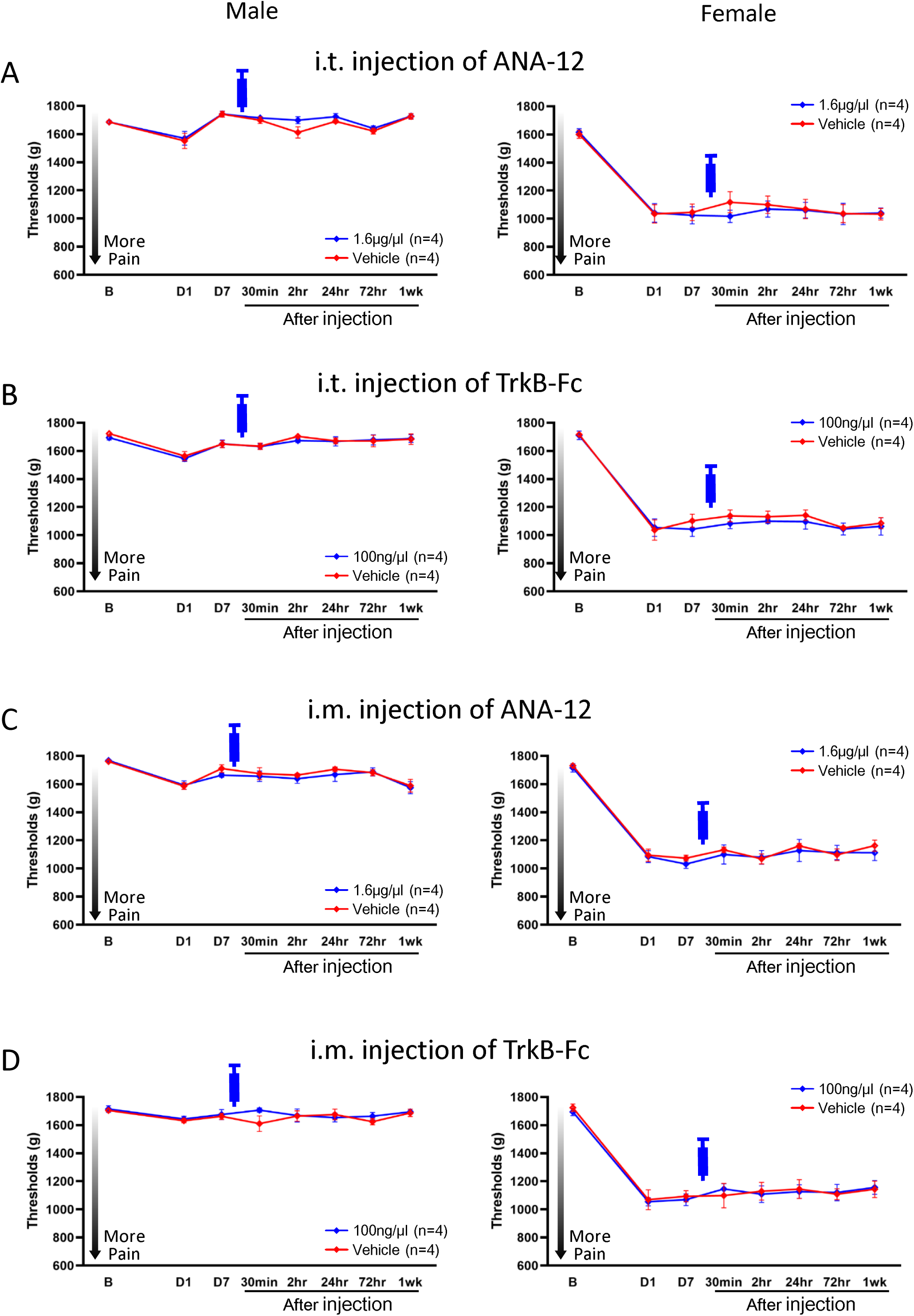
Contralateral muscle withdrawal thresholds with pharmacological inhibition intrathecally or intramuscularly 1wk after induction of the model. BDNF inhibitors intrathecally or intramuscularly 1wk after induction of the model had no effect on muscle withdrawal thresholds on contralateral side in male or female mice (A–D). Data is reported as mean ± S.E.M. *, versus vehicle control; p<0.05. i.m., intramuscularly; i.t., intrathecally.

**Supplemental Fig. 5.**
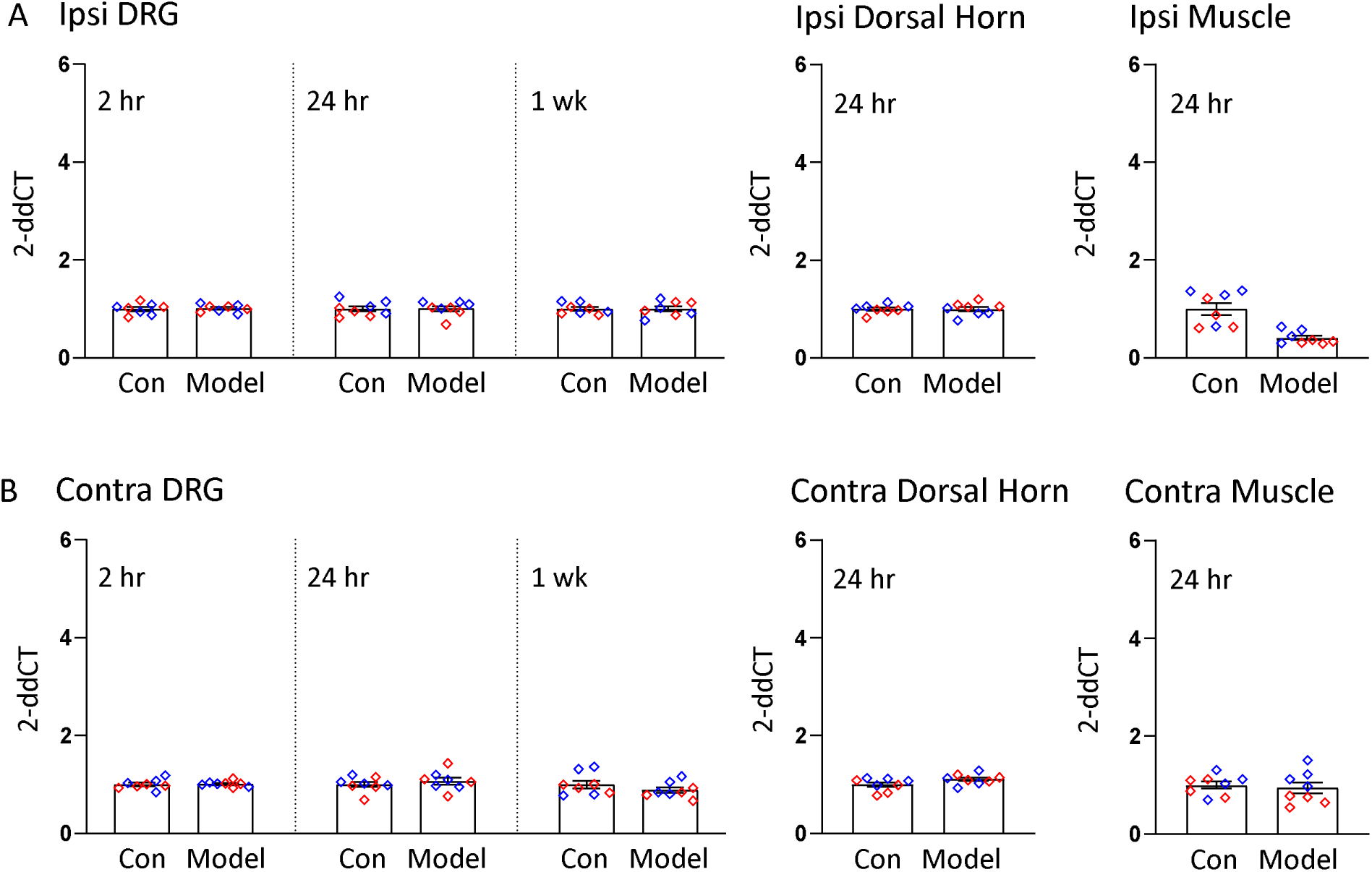
TrkB mRNA expressions after induction of the activity-induced muscle pain. TrkB mRNA showed no difference in either ipsilateral or contralateral L4–L6 DRG, spinal dorsal horn, or gastrocnemius muscle 24hr after induction of the model, when compared to pain free controls (A, B). Blue plots represent male data, Red plots represent female data. Data is reported as mean ± S.E.M. N=8 in each group. *, p<0.05. DRG, dorsal root ganglion; TrkB, tropomyosin receptor kinase B; mRNA, messenger ribonucleic acid.

**Supplemental Fig. 6.**
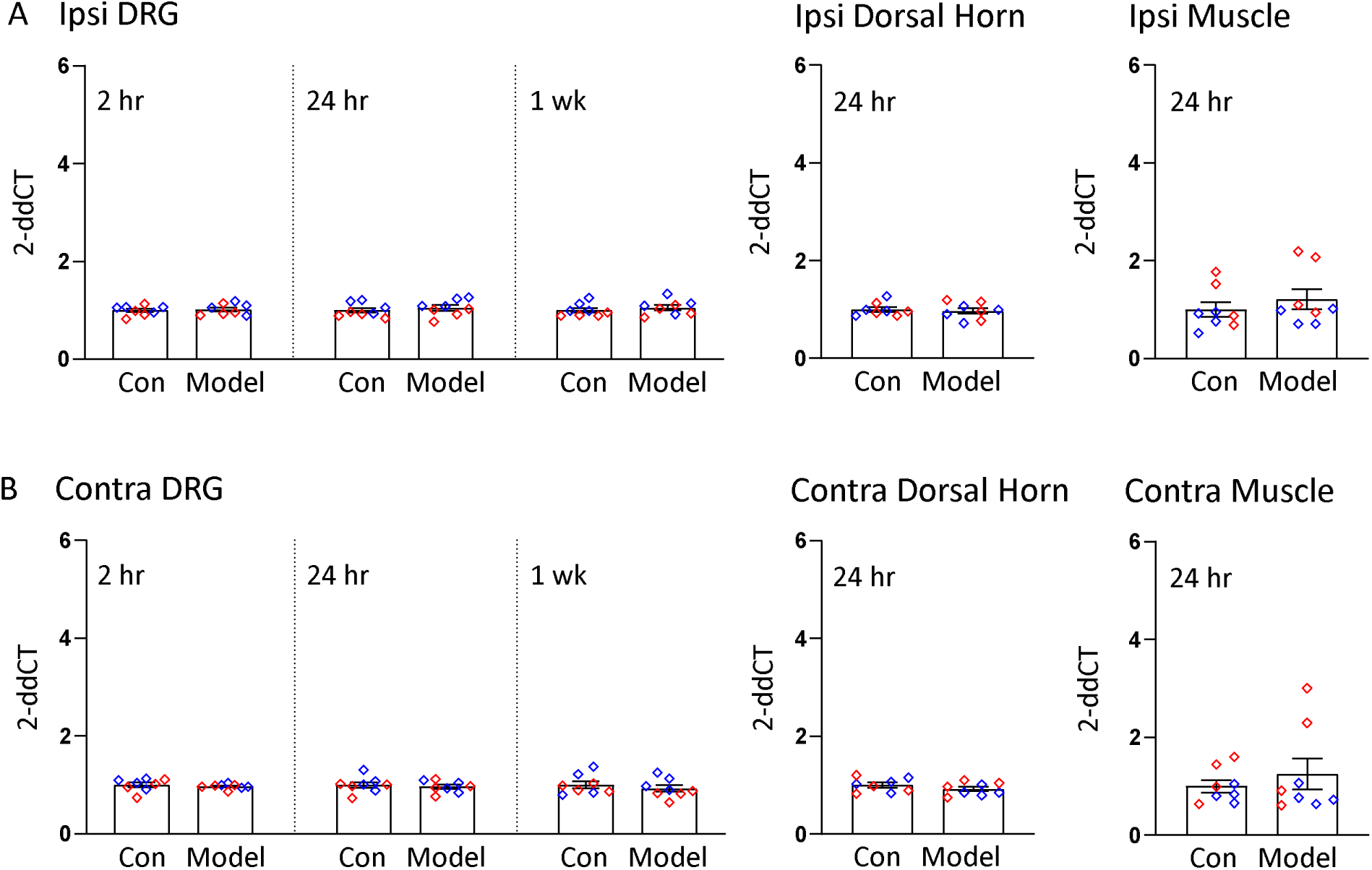
p75^NTR^ mRNA expressions after induction of the activity-induced muscle pain. p75^NTR^ mRNA showed no difference in either ipsilateral or contralateral L4–L6 DRG, spinal dorsal horn, or gastrocnemius muscle 24hr after induction of the model, when compared to pain free controls (A, B). Blue plots represent male data, Red plots represent female data. Data is reported as mean ± S.E.M. N=8 in each group. *, p<0.05. DRG, dorsal root ganglion; mRNA, messenger ribonucleic acid; p75^NTR^, p75 neurotrophin receptor.

**Supplemental Fig. 7.**
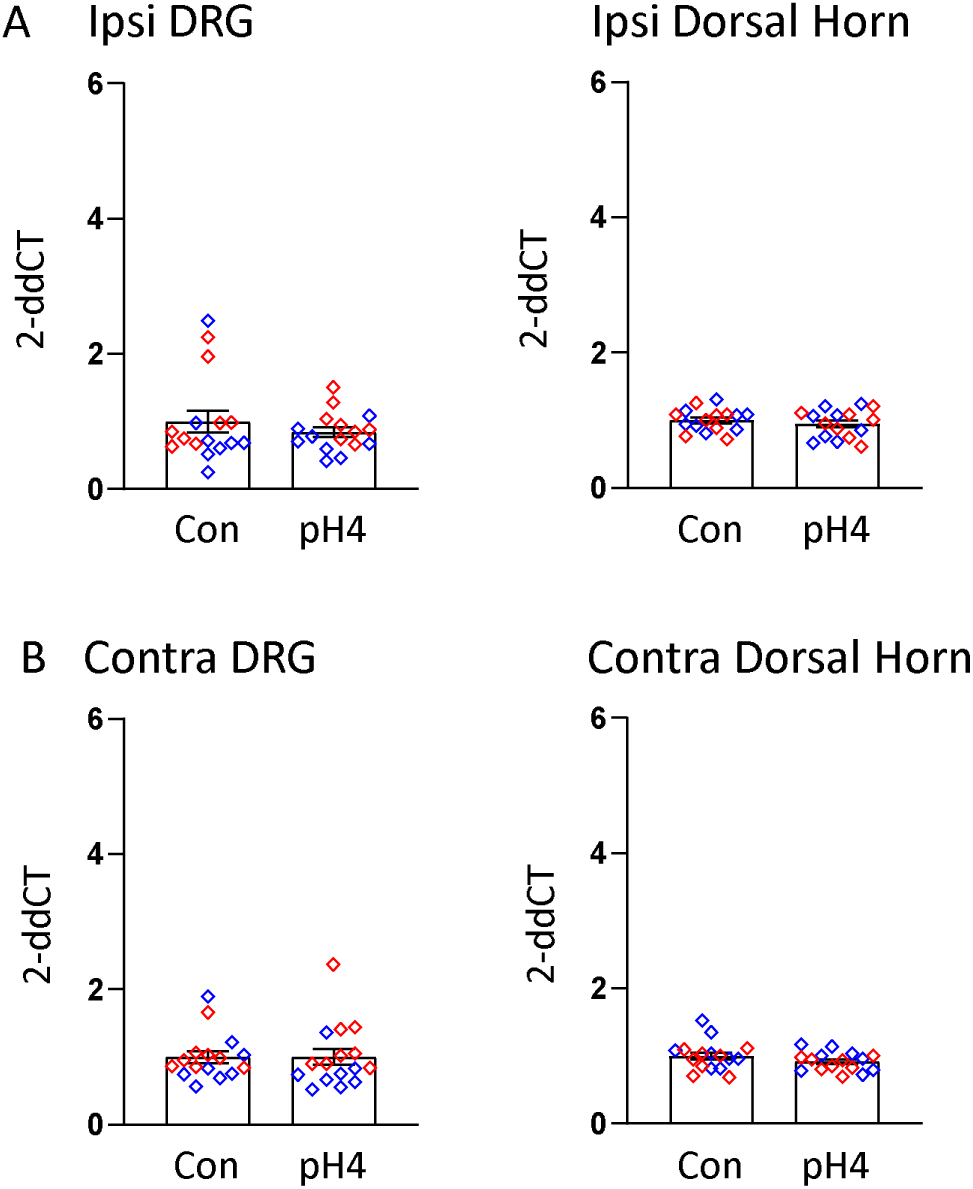
BDNF mRNA expressions in the animal model of pH4 pain. BDNF mRNA showed no difference in either ipsilateral or contralateral L4–L6 DRG, or spinal dorsal horn 24hr after induction of the model, when compared to pain free controls (A, B). Blue plots represent male data, Red plots represent female data. Data is reported as mean ± S.E.M. N=16 in each group. *, p<0.05. BDNF, brain-derived neurotrophic factor; DRG, dorsal root ganglion; mRNA, messenger ribonucleic acid.

## References

1. Institute of Medicine (US) Committee on Advancing Pain Research, Care, and Education. Relieving Pain in America: A Blueprint for Transforming Prevention, Care, Education, and Research. Washington (DC): National Academies Press (US); 2011.

2. Dahlhamer J LJ, Zelaya C, Nahin R, Mackey S, DeBar L, Kerns R, Von Korff M, Porter L, Helmick C. Prevalence of Chronic Pain and High-Impact Chronic Pain Among Adults – United States, 2016. MMWR Morb Mortal Wkly Rep. 2018; 67:1001–1006.

3. GBD 2019 Diseases and Injuries Collaborators. Global burden of 369 diseases and injuries in 204 countries and territories, 1990-2019: a systematic analysis for the Global Burden of Disease Study 2019. Lancet. 2020; 396:1204–1222. Erratum in: Lancet. 2020; 396:1562.

4. Kosek E, Ekholm J, Hansson P. Modulation of pressure pain thresholds during and following isometric contraction in patients with fibromyalgia and in healthy controls. Pain. 1996; 64:415–423.

5. Landmark T, Romundstad P, Borchgrevink PC, Kaasa S, Dale O. Associations between recreational exercise and chronic pain in the general population: Evidence from the HUNT 3 study. Pain. 2011; 152:2241–2247.

6. Landmark T, Romundstad PR, Borchgrevink PC, Kaasa S, Dale O. Longitudinal Associations between Exercise and Pain in the General Population - The HUNT Pain Study. Plos One. 2013; 8:e65279.

7. Gregory NS, Gibson-Corley K, Frey-Law L, Sluka KA. Fatigue-enhanced hyperalgesia in response to muscle insult: induction and development occur in a sex-dependent manner. Pain. 2013; 154:2668–2676.

8. Gregory N, Brito R, Fusaro MCG, Sluka KA. ASIC3 Is Required for Development of Fatigue-Induced Hyperalgesia. Mol Neurobiol. 2016; 53:1020–1030.

9. Oliveira-Fusaro MC, Gregory NS, Kolker SJ, Rasmussen L, Allen LAH, Sluka KA. P2X4 Receptors on Muscle Macrophages Are Required for Development of Hyperalgesia in an Animal Model of Activity-Induced Muscle Pain. Mol Neurobiol. 2020; 57:1917–1929.

10. Hayashi K, Lesnak JB, Plumb AN, Rasmussen LA, Sluka KA. P2X7-NLRP3-Caspase-1 signaling mediates activity-induced muscle pain in male but not female mice. Pain. 2023; 164:1860–1873.

11. Ren K, Torres R. Role of interleukin-1beta during pain and inflammation. Brain Res Rev. 2009; 60:57–64.

12. Starobova H, Alshammari A, Winkler IG, Vetter I. The role of the neuronal microenvironment in sensory function and pain pathophysiology. J Neurochem. 2022. In press.

13. Lever IJ, Bradbury EJ, Cunningham JR, Adelson DW, Jones MG, McMahon SB, Marvizón JC, Malcangio M. Brain-derived neurotrophic factor is released in the dorsal horn by distinctive patterns of afferent fiber stimulation. J Neurosci. 2001; 21:4469–4477.

14. Pezet S, McMahon SB. Neurotrophins: mediators and modulators of pain. Annu Rev Neurosci. 2006; 29:507–538.

15. Cappoli N, Tabolacci E, Aceto P, Dello Russo C. The emerging role of the BDNF-TrkB signaling pathway in the modulation of pain perception. J Neuroimmunol. 2020; 349:577406.

16. Thompson SW, Bennett DL, Kerr BJ, Bradbury EJ, McMahon SB. Brain-derived neurotrophic factor is an endogenous modulator of nociceptive responses in the spinal cord. Proc Natl Acad Sci U S A. 1999; 96:7714–7718.

17. Michael GJ, Averill S, Nitkunan A, Rattray M, Bennett DL, Yan Q, Priestley JV. Nerve growth factor treatment increases brain-derived neurotrophic factor selectively in TrkA-expressing dorsal root ganglion cells and in their central terminations within the spinal cord. J Neurosci. 1997; 17:8476–8490.

18. Cho HJ, Increased brain-derived neurotrophic factor immunoreactivity in rat dorsal root ganglia and spinal cord following peripheral inflammation. Brain Res. 1997; 764:269–272.

19. Chen W, Walwyn W, Ennes HS, Kim H, McRoberts JA, Marvizón JC. BDNF released during neuropathic pain potentiates NMDA receptors in primary afferent terminals. Eur J Neurosci. 2014; 39:1439–1454.

20. Kohara K, Kitamura A, Morishima M, Tsumoto T. Activity-dependent transfer of brain-derived neurotrophic factor to postsynaptic neurons. Science. 2001; 291:2419–2423.

21. Walker SM, Mitchell VA, White DM, Rush RA, Duggan AW. Release of immunoreactive brain-derived neurotrophic factor in the spinal cord of the rat following sciatic nerve transection. Brain Res. 2001; 899:240–247.

22. Zhou XF, Rush RA. Endogenous brain-derived neurotrophic factor is anterogradely transported in primary sensory neurons. Neuroscience. 1996; 74:945–953.

23. Merighi A, Salio C, Ghirri A, Lossi L, Ferrini F, Betelli C, Bardoni R. BDNF as a pain modulator. Prog Neurobiol. 2008; 85:297–317.

24. Malcangio M, Lessmann V. A common thread for pain and memory synapses? Brain-derived neurotrophic factor and trkB receptors. Trends Pharmacol Sci. 2003; 24:116–121.

25. Mapplebeck JCS, Lorenzo LE, Lee KY, Gauthier C, Muley MM, De Koninck Y, Prescott SA, Salter MW. Chloride Dysregulation through Downregulation of KCC2 Mediates Neuropathic Pain in Both Sexes. Cell Rep. 2019; 28:590–596.e4.

26. Gómez-Pinilla F, Ying Z, Roy RR, Molteni R, Edgerton VR. Voluntary exercise induces a BDNF-mediated mechanism that promotes neuroplasticity. J Neurophysiol. 2002; 88:2187–2195.

27. Luo C, Zhong XL, Zhou FH, Li JY, Zhou P, Xu JM, Song B, Li CQ, Zhou XF, Dai RP. Peripheral Brain Derived Neurotrophic Factor Precursor Regulates Pain as an Inflammatory Mediator. Sci Rep. 2016; 6:27171.

28. Gowler PRW, Li L, Woodhams SG, Bennett AJ, Suzuki R, Walsh DA, Chapman V. Peripheral brain-derived neurotrophic factor contributes to chronic osteoarthritis joint pain. Pain. 2020; 161:61–73.

29. Zhao J, Seereeram A, Nassar MA, Levato A, Pezet S, Hathaway G, Morenilla-Palao C, Stirling C, Fitzgerald M, McMahon SB, Rios M, Wood JN; London Pain Consortium. Nociceptor-derived brain-derived neurotrophic factor regulates acute and inflammatory but not neuropathic pain. Mol Cell Neurosci. 2006; 31:539–548.

30. Dembo T, Braz JM, Hamel KA, Kuhn JA, Basbaum AI. Primary Afferent-Derived BDNF Contributes Minimally to the Processing of Pain and Itch. eNeuro. 2018; 5:ENEURO.0402-18.2018.

31. Sikandar S, Minett MS, Millet Q, Santana-Varela S, Lau J, Wood JN, Zhao J. Brain-derived neurotrophic factor derived from sensory neurons plays a critical role in chronic pain. Brain. 2018; 141:1028–1039.

32. Sluka KA, Kalra A, Moore SA. Unilateral intramuscular injections of acidic saline produce a bilateral, long-lasting hyperalgesia. Muscle Nerve. 2001; 24:37–46.

33. Sahlin K, Harris RC, Nylind B, Hultman E. Lactate Content and Ph in Muscle Samples Obtained after Dynamic Exercise. Pflug Arch Eur J Phy. 1976; 367:143–149.

34. Spriet LL, Soderlund K, Thomson JA, Hultman E. Ph Measurement in Human Skeletal-Muscle Samples - Effect of Phosphagen Hydrolysis. J Appl Physiol. 1986; 61:1949–1954.

35. Victor RG, Bertocci LA, Pryor SL, Nunnally RL. Sympathetic-Nerve Discharge Is Coupled to Muscle-Cell Ph during Exercise in Humans. J Clin Invest. 1988; 82:1301–1305.

36. Skyba DA, Radhakrishnan R, Sluka KA. Characterization of a method for measuring primary hyperalgesia of deep somatic tissue. J Pain. 2005; 6:41–47.

37. Fan F, Tang Y, Dai H, Cao Y, Sun P, Chen Y, Chen A, Lin C. Blockade of BDNF signalling attenuates chronic visceral hypersensitivity in an IBS-like rat model. Eur J Pain. 2020; 24:839–850.

38. Melemedjian OK, Tillu DV, Asiedu MN, Mandell EK, Moy JK, Blute VM, Taylor CJ, Ghosh S, Price TJ. BDNF regulates atypical PKC at spinal synapses to initiate and maintain a centralized chronic pain state. Mol Pain. 2013; 9:12.

39. Moy JK, Szabo-Pardi T, Tillu DV, Megat S, Pradhan G, Kume M, Asiedu MN, Burton MD, Dussor G, Price TJ. Temporal and sex differences in the role of BDNF/TrkB signaling in hyperalgesic priming in mice and rats. Neurobiol Pain. 2018; 5:100024.

40. Lee-Kubli CAG, Calcutt NA. Altered rate-dependent depression of the spinal H-reflex as an indicator of spinal disinhibition in models of neuropathic pain. Pain. 2014; 155:250–260.

41. Ruscheweyh R, Forsthuber L, Schoffnegger D, Sandkühler J. Modification of classical neurochemical markers in identified primary afferent neurons with Abeta-, Adelta-, and C-fibers after chronic constriction injury in mice. J Comp Neurol. 2007; 502:325–336.

42. Lesnak JB, Nakhla DS, Plumb AN, McMillan A, Saha S, Gupta N, Xu Y, Phruttiwanichakun P, Rasmussen L, Meyerholz DK, Salem AK, Sluka KA. Selective androgen receptor modulator microparticle formulation reverses muscle hyperalgesia in a mouse model of widespread muscle pain. Pain. 2023; 164:1512–1523.

43. Light AR, Hughen RW, Zhang J, Rainier J, Liu ZQ, Lee J. Dorsal root ganglion neurons innervating skeletal muscle respond to physiological combinations of protons, ATP, and lactate mediated by ASIC, P2X, and TRPV1. J Neurophysiol. 2008; 100:1184–1201.

44. Lin YT, Ro LS, Wang HL, Chen JC. Up-regulation of dorsal root ganglia BDNF and trkB receptor in inflammatory pain: an in vivo and in vitro study. J Neuroinflammation. 2011; 8:126.

45. Chen WK, Liu IY, Chang YT, Chen YC, Chen CC, Yen CT, Shin HS, Chen CC. Ca(v)3.2 T-Type Ca2+ Channel-Dependent Activation of ERK in Paraventricular Thalamus Modulates Acid-Induced Chronic Muscle Pain. J Neurosci. 2010; 30:10360–10368.

46. Hoeger-Bement MK, Sluka KA. Phosphorylation of CREB and mechanical hyperalgesia is reversed by blockade of the cAMP pathway in a time-dependent manner after repeated intramuscular acid injections. J Neurosci. 2003; 23:5437–5445.

47. Spandidos A, Wang XW, Wang HJ, Seed B. PrimerBank: a resource of human and mouse PCR primer pairs for gene expression detection and quantification. Nucleic Acids Res. 2010; 38:D792–D799.

48. Thornton B, Basu C. Real-Time PCR (qPCR) Primer Design Using Free Online Software. Biochem Mol Biol Edu. 2011; 39:145–154.

49. Wang XW, Spandidos A, Wang HJ, Seed B. PrimerBank: a PCR primer database for quantitative gene expression analysis, 2012 update. Nucleic Acids Res. 2012; 40:D1144–D1149.

50. Schmittgen TD, Livak KJ. Analyzing real-time PCR data by the comparative C-T method. Nat Protoc. 2008; 3:1101–1108.

51. Lesnak JB, Fahrion A, Helton A, Rasmussen L, Andrew M, Cunard S, Huey M, Kreber A, Landon J, Siwiec T, Todd K, Frey-Law LA, Sluka KA. Resistance training protects against muscle pain through activation of androgen receptors in male and female mice. Pain. 2022; 163:1879–1891.

52. Sahlin K, Harris RC, Nylind B, Hultman E. Lactate Content and Ph in Muscle Samples Obtained after Dynamic Exercise. Pflug Arch Eur J Phy. 1976; 367:143–149.

53. Spriet LL, Soderlund K, Thomson JA, Hultman E. Ph Measurement in Human Skeletal-Muscle Samples - Effect of Phosphagen Hydrolysis. J Appl Physiol. 1986; 61:1949–1954.

54. Victor RG, Bertocci LA, Pryor SL, Nunnally RL. Sympathetic-Nerve Discharge Is Coupled to Muscle-Cell Ph during Exercise in Humans. J Clin Invest. 1988; 82:1301–1305.

55. Gregory NS, Gautam M, Benson CJ, Sluka KA. Acid Sensing Ion Channel 1a (ASIC1a) Mediates Activity-induced Pain by Modulation of Heteromeric ASIC Channel Kinetics. Neuroscience. 2018; 386:166–174.

56. Immke DC, McCleskey EW. Lactate enhances the acid-sensing Na+ channel on ischemia-sensing neurons. Nat Neurosci. 2001; 4:869–870.

57. Birdsong WT, Fierro L, Williams FG, Spelta V, Naves LA, Knowles M, Marsh-Haffner J, Adelman JP, Almers W, Elde RP, McCleskey EW. Sensing Muscle Ischemia: Coincident Detection of Acid and ATP via Interplay of Two Ion Channels. Neuron. 2010; 68:739–749.

58. Gregory NS, Whitley PE, Sluka KA. Effect of Intramuscular Protons, Lactate, and ATP on Muscle Hyperalgesia in Rats. PLoS One. 2015; 10:e0138576.

59. Ross JL, Queme LF, Shank AT, Hudgins RC, Jankowski MP. Sensitization of group III and IV muscle afferents in the mouse after ischemia and reperfusion injury. J Pain. 2014; 15:1257–1270.

60. Ross JL, Queme LF, Cohen ER, Green KJ, Lu P, Shank AT, An S, Hudgins RC, Jankowski MP. Muscle IL1β Drives Ischemic Myalgia via ASIC3-Mediated Sensory Neuron Sensitization. J Neurosci. 2016; 36:6857–6871.

61. Noma N, Shinoda M, Honda K, Kiyomoto M, Dezawa K, Nakaya Y, Komiyama O, Imamura Y, Iwata K. Interaction of IL-1β and P2X(3) receptor in pathologic masseter muscle pain. J Dent Res. 2013; 92:456–460.

62. Sluka KA, Price MP, Breese NM, Stucky CL, Wemmie JA, Welsh MJ. Chronic hyperalgesia induced by repeated acid injections in muscle is abolished by the loss of ASIC3, but not ASIC1. Pain. 2003; 106:229–239.

63. Gong WY, Abdelhamid RE, Carvalho CS, Sluka KA. Resident Macrophages in Muscle Contribute to Development of Hyperalgesia in a Mouse Model of Noninflammatory Muscle Pain. J Pain. 2016; 17:1081–1094.

64. Ferreira SH, Lorenzetti BB, Bristow AF, Poole S. Interleukin-1 beta as a potent hyperalgesic agent antagonized by a tripeptide analogue. Nature. 1988; 334:698–700.

65. Lesnak JB, Berardi G, Sluka KA. Influence of routine exercise on the peripheral immune system to prevent and alleviate pain. Neurobiol Pain. 2023; 13:100126.

66. Winston JH, Li Q, Sarna SK. Chronic prenatal stress epigenetically modifies spinal cord BDNF expression to induce sex-specific visceral hypersensitivity in offspring. Neurogastroenterol Motil. 2014; 26:715–730.

67. Sorge RE, Mapplebeck JC, Rosen S, Beggs S, Taves S, Alexander JK, Martin LJ, Austin JS, Sotocinal SG, Chen D, Yang M, Shi XQ, Huang H, Pillon NJ, Bilan PJ, Tu Y, Klip A, Ji RR, Zhang J, Salter MW, Mogil JS. Different immune cells mediate mechanical pain hypersensitivity in male and female mice. Nat Neurosci. 2015; 18:1081–1083.

68. Burgos-Vega CC, Quigley LD, Avona A, Price T, Dussor G. Dural stimulation in rats causes brain-derived neurotrophic factor-dependent priming to subthreshold stimuli including a migraine trigger. Pain. 2016; 157:2722–2730.

69. Zhou LJ, Peng J, Xu YN, Zeng WJ, Zhang J, Wei X, Mai CL, Lin ZJ, Liu Y, Murugan M, Eyo UB, Umpierre AD, Xin WJ, Chen T, Li M, Wang H, Richardson JR, Tan Z, Liu XG, Wu LJ. Microglia are indispensable for synaptic plasticity in the spinal dorsal horn and p. Cell Rep. 2019; 27:3844–3859.e6.

70. Dedek A, Xu J, Lorenzo LÉ, Godin AG, Kandegedara CM, Glavina G, Landrigan JA, Lombroso PJ, De Koninck Y, Tsai EC, Hildebrand ME. Sexual dimorphism in a neuronal mechanism of spinal hyperexcitability across rodent and human models of pathological pain. Brain. 2022; 145:1124–1138.

71. Arévalo JC, Deogracias R. Mechanisms Controlling the Expression and Secretion of BDNF. Biomolecules. 2023; 13:789.

72. Lessmann V, Brigadski T. Mechanisms, locations, and kinetics of synaptic BDNF secretion: an update. Neurosci Res. 2009; 65:11–22. Erratum in: Neurosci Res. 2009; 65:316–317.

73. Ohtori S, Takahashi K, Moriya H. Inflammatory pain mediated by a phenotypic switch in brain-derived neurotrophic factor-immunoreactive dorsal root ganglion neurons innervating the lumbar facet joints in rats. Neurosci Lett. 2002; 323:129–132.

74. Zhou XF, Chie ET, Deng YS, Zhong JH, Xue Q, Rush RA, Xian CJ. Injured primary sensory neurons switch phenotype for brain-derived neurotrophic factor in the rat. Neuroscience. 1999; 92:841–853.

75. Ha SO, Kim JK, Hong HS, Kim DS, Cho HJ. Expression of brain-derived neurotrophic factor in rat dorsal root ganglia, spinal cord and gracile nuclei in experimental models of neuropathic pain. Neuroscience. 2001; 107: 301–309.

76. Plumb AN, Lesnak JB, Berardi G, Hayashi K, Janowski AJ, Smith AF, Bailey D, Kerkman C, Kienenberger Z, Martin B, Patterson E, Van Roekel H, Vance CGT, Sluka KA. Standing on the shoulders of bias: lack of transparency and reporting of critical rigor characteristics in pain research. Pain. 2023; 164:1775–1782.

77. Hankerd K, Koo H, McDonough KE, Wang J, Pariyar R, Tang SJ, Chung JM, La JH. Gonadal hormone-dependent nociceptor sensitization maintains nociplastic pain state in female mice. Pain. 2023; 164:402–412.

78. Lesnak JB, Inoue S, Lima L, Rasmussen L, Sluka KA. Testosterone protects against the development of widespread muscle pain in mice. Pain. 2020; 161:2898–2908.

79. Lesnak JB, Hayashi K, Plumb AN, Janowski AJ, Chimenti MS, Sluka KA. The impact of sex and physical activity on the local immune response to muscle pain. Brain Behav Immun. 2023; 111:4–20.

